# DNA-based ForceChrono Probes for Deciphering Single-Molecule Force Dynamics in Living Cells

**DOI:** 10.1101/2024.03.09.584267

**Authors:** Yuru Hu, Hongyun Li, Chen Zhang, Jingjing Feng, Wenxu Wang, Wei Chen, Miao Yu, Xinghua Zhang, Zheng Liu

## Abstract

Accurate measurement of mechanical forces in cells is key to understanding how cells sense and respond to mechanical stimuli, a central aspect of mechanobiology. However, accurately quantifying dynamic forces at the single-molecule level in living cells is a significant challenge. Here, we’ve developed the DNA-based ForceChrono probe to enable in-depth studies of integrin force dynamics at the single-molecule level in living cells. By illuminating two distinct mechanical points and circumventing the inherent fluctuations of single-molecule fluorescence, the ForceChrono probe enables analysis of the complex dynamics of mechanical forces at the single-molecule level, such as loading rates and durations. Our results refine previous broad estimates of cellular loading rates to a more precise range of 0.5 to 2 pN/s, shedding light on the specifics of cellular mechanics. In addition, this study reveals a critical link between the magnitude and duration of integrin forces, consistent with the catch-bond behavior demonstrated in vitro. The ForceChrono probe has distinct advantages, such as precise analysis of single-molecule force dynamics and robust resistance to fluorescence fluctuations, which will significantly advance our understanding of cell adhesion and mechanotransduction at the single-molecule level.

## Introduction

The cellular microenvironment is not just a passive milieu where cells exist, but rather a dynamic theater where intricate molecular dramas unfold. Cells are not mere spectators; they actively interpret and respond to the myriad of mechanical cues presented to them^1^. These cues, often manifested as forces exerted on cellular receptors, play a crucial role in dictating cellular behavior^2,3^. To fully understand this complex interplay, it’s essential to delve deeper into the three pivotal parameters that characterize these forces: the magnitude, the loading rate, and the sustained duration^4,5^.

Over the past decade, various types of immobilized molecular fluorescent tension probes have emerged as a revolutionary tool in mechanobiology, providing a detailed lens through which the magnitude of cellular forces can be observed and quantified at the molecular level^6–12^. Notable examples include the elucidation of integrin-mediated force transduction^8,10,13,14^, the dynamics of T-cell receptor (TCR) forces^15,16^, and the mechanisms underlying cellular rigidity sensing^17^. These studies have not only quantified the magnitude characteristics of cellular forces but have also underscored their pivotal role in driving the mechanisms of mechanical force transduction. While the achievements in quantifying force magnitude are commendable, the field still grapples with challenges in measuring other critical parameters, notably the force loading rate and force duration.

The duration of force reveals how long a protein or a molecular complex can resist a mechanical force before detaching or undergoing a conformational change^18–21^. This parameter is vital for understanding the mechanical stability and resilience of key molecular players in mechanotransduction pathways^18,22,23^. Moreover, the duration of these forces can significantly influence molecular interactions^24^. The loading rate can determine how proteins unfold, activate, or even interact with other molecules, thereby dictating the immediate cellular response to mechanical stimuli^22,23,25–28^. A key aspect to highlight in the study of cellular mechanotransduction is the significant dependence of rupture force and bonding lifetime on the loading rate^20,29^. This dependency underscores the complexity of understanding cellular mechanics, as our current knowledge of physiologically relevant cellular loading rates is limited. Often, studies resort to assuming an arbitrary loading rate or reporting a spectrum of rates observed throughout the research. The estimated range of force loading rates exerted by living cells, as predicted by various research groups, is remarkably broad, spanning from a minimal 0.007 pN/s to an extreme of 10^7^ pN/s^27,30,31^. This wide range often leads researchers to assume arbitrary loading rates or report a series of rates to explain observed phenomena, creating confusion in understanding cellular mechanics. Understanding and quantifying these parameters in living cells is not just a biophysical curiosity but a necessity to unravel the intricate dance of molecules under mechanical stress. Here, we designed a ForceChrono probe that combined two DNA hairpin probes with distinct force thresholds and fluorophores, enabling a tiered system for force measurement. This design overcomes the limitations of traditional DNA hairpin-based tension probes that only report force threshold values. This multi-tiered approach not only measures information about the magnitude of the single-molecule force in living cells, but also offers insights into the dynamics of the force application, both in terms of the loading rate and duration of the force. Our results using the ForceChrono probe reveal a correlation between the magnitude and duration of integrin forces and accurately measure the loading rate of cells in different environments, as well as the relevant influences on these values. These studies fill a critical gap in current research methods and thus deepen our understanding of mechanotransduction pathways. Overall, we demonstrate that the DNA-based ForceChrono probe is uniquely suited to reveal the complex dynamics of integrin force at the single-molecule level, providing new insights into the molecular mechanisms of cellular mechanotransduction.

## Results

### Development of ForceChrono Probes

The ForceChrono probes comprise a dual DNA hairpin system, with each hairpin tagged with a unique fluorophore—Cy3B for the lower force threshold and Atto647N for the higher (Figure 1A and S1A). These probes are synthesized and assembled using a modular strategy (Figure 1A and S1B). By adjusting the force application geometry of the DNA hairpin, GC content, stem lengths, or loop size (Table S1-3), these hairpins are engineered to unfold sequentially at specific mechanical force thresholds, enabling real-time measurement of force loading rates. A fluorescence quenching moiety, BHQ2, is strategically positioned at the hairpin’s junction, quenching both dyes in the “off” state. To further enhance optical sensitivity and prevent photobleaching, the entire probe is immobilized on a 3.5 nm gold nanoparticle. This design not only provides an additional layer of fluorescence quenching through Nanoscale Gold Surface Resonance Energy Transfer (NEST)^10,15,32^ but also ensures the reliability and accuracy of ON-OFF signals in subsequent single-molecule fluorescence imaging experiments.

**Figure 1.**
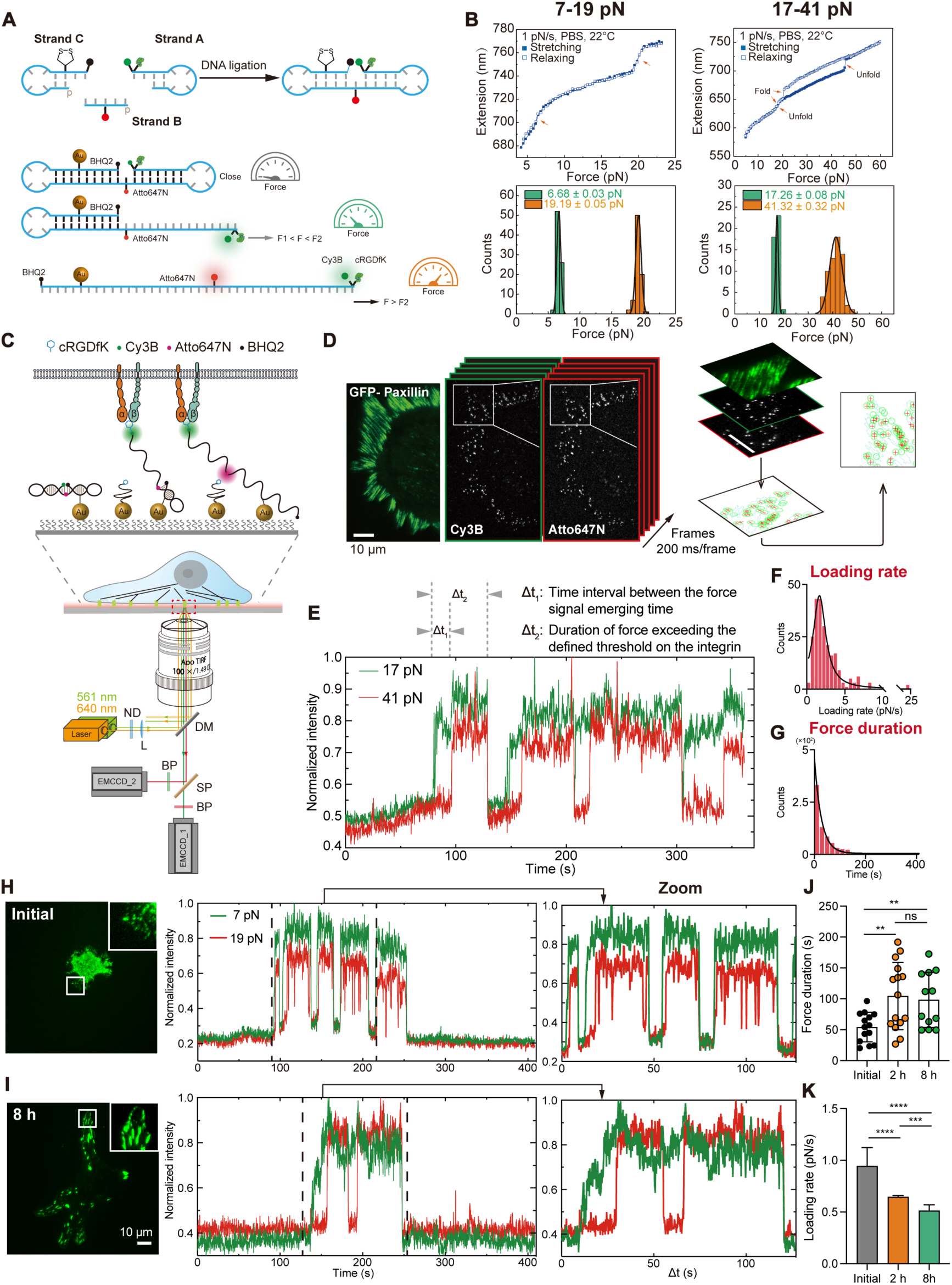
Development of ForceChrono probes for visualizing individual integrin mechanical force dynamics in living cells. **A,** Schematic of the ForceChrono probes assembly (upper) and structure (lower). **B,** Calibration curves of ForceChrono probes with SMMTs. Upper: representative force-extension curves of 7-19 pN and 17-41 pN probes. Lower: Histograms and Gaussian-fitting results of applied forces at the unfolding force events. **C,** Microscope setup for simultaneous single-molecule fluorescence imaging of two channels. **D,** Representative fluorescence images of paxillin-GFP, Cy3B and Atto647N from the TIRF microscope. Integrin force signals in the Cy3B and Atto647N channels were imaged using a 200 ms exposure time for 2000 frames without intervals. Then the images were calibrated and the localization information of single molecules was extracted. **E,** Representative fluorescence trace of individual integrin force signals reported by 17-41 pN. Δt1 represents the time interval between the emergence of the force signal. Δt2 represents the duration of the mechanical force on the integrin-ligand bond exceeding the unfolding force. **F,** Distribution of integrin mechanical force loading rate and the fitting result of Lorentzian Distribution. **G,** Distribution of integrin mechanical force duration and the fitting result of exponential decay. **H, I,** Left: Representative paxillin-GFP images of MEF cells spreading out at different times. Right: Representative fluorescence trace of integrin force signals reported by 7-19 pN. **J,** Average duration of force on integrins measured using 7-19 pN probes at different cell spreading times. Each point represents the mean duration of a cell. The error bars represent mean ± s.d. from three independent experiments. **K,** Average loading rate of force on integrins measured using 7-19 pN probes at different cell spreading times. The error bars represent mean ± 95% CI from three independent experiments. Two-tailed Student’s *t*-tests are used to assess statistical significance. Scale bar=10 μm.

Before cellular application, the ForceChrono probes were rigorously calibrated using single-molecule magnetic tweezers (SMMTs), confirming that the mechanical thresholds for hairpin unfolding aligned with theoretical predictions (Figure S2 and Table S4). Two versions were developed to cover a broader mechanical range (Figure 1B): one sensitive to 7 pN and 19 pN forces, and another to 17 pN and 41 pN. The probes were diluted to single-molecule concentrations and covalently attached to gold nanoparticles, along with non-fluorescent cRGDfK peptides for cellular experiments. This created a functionalized substrate for Mouse Embryonic Fibroblast (MEF) cells (Figure S1).

Imaging was conducted using Total Internal Reflection Fluorescence (TIRF) microscopy (Figure 1C), equipped with dual EMCCDs and a dichroic short-wavelength pass filter. This setup allowed for the simultaneous, real-time capture of both Cy3B and Atto647N emissions. High temporal resolution was maintained with an exposure time of 200 ms across 2000 continuous frames (Figure 1D and Movie S1). The stability of the fluorescent dyes was validated through photobleaching experiments (Figure S3A-D). We calibrated the acquired images and extracted the single-molecule signal to generate time-resolved fluorescence trajectories, yielding both the loading rate and duration of integrin forces (Figure 1E). In particular, the time difference between the sequential activations of the two fluorophores directly measures the force loading rate, given the known mechanical thresholds of the hairpins (Figure 1F). Furthermore, the duration for which both fluorophores remain activated indicates sustained mechanical force, offering an unprecedented measure of force duration at the single-molecule level. Specifically, the first hairpin’s design in unzipping mode eliminates hysteresis between unfolding and refolding forces (Figure 1B), meaning the duration for which the first fluorescent molecule (Cy3B) is illuminated should indeed represent the force duration (Figure 1G). Additionally, we used a probe with only one hairpin structure as a control and found that the opening of the second hairpin does not impact the measurement of force duration (Figure S3E-G). This ensures the accuracy of our measurements and validates the comparability of the loading rate results across different probes.

In nearly all single-molecule events, both fluorescent molecules on the ForceChrono probe continued to glow simultaneously for a duration before extinguishing simultaneously (Figure 1E). This suggests that the force on the integrin gradually increases and sustains for a period before abruptly terminating. This abrupt termination is likely due to either a break in the integrin-ligand bond outside the cell membrane or an internal protein-protein bond break, such as talin-integrin^33^. This contrasts with a slow relaxation scenario, where the two fluorescent molecules would extinguish sequentially due to the different refolding forces of the hairpins.

To further explore the temporal dynamics of single integrin force, we employed MEFs stably expressing Paxillin-GFP and spread them onto substrates modified with 7-19 pN ForceChrono probes (Figure 1H-K). During the early stages of adhesion, cells formed highly dynamic needle-like and dot-like adhesion structures (Figure 1H). At this stage, the duration of force on integrin was short, averaging around 37 seconds, and the loading rate was high, approximately 0.9 pN/s. However, as the cells spread over 2 hours, both the force duration and the loading rate underwent significant changes (Figure 1J and 1K). By 8 hours of spreading (Figure 1I), the cells exhibited obvious polarization, and the loading rate had decreased to about 0.5 pN/s.

These observations indicate an apparent spatio-temporal heterogeneity in the mechanical dynamics of integrins at different stages of cell spreading. The changes in mechanical dynamics may be closely related to the stability of the adhesion structures and the slowing down of actin retrograde flow^30^. This spatio-temporal heterogeneity underscores the complexity of mechanical force regulation in cellular processes and highlights the utility of ForceChrono probes in capturing these dynamics at the single-molecule level.

### Probing the force duration of a single integrin in living cells

When dissecting the intricate dynamics of integrin-ligand interactions within living cells using ForceChrono probes, it is essential to evaluate the sensitivity of the probes (Figure 2A). Specifically, it is crucial to determine whether the hairpin structures of the probe can open when the mechanical force on the integrin reaches a certain threshold, with the recovery of the fluorescence signal. To address this, we designed a ForceChrono probe with nearly identical threshold forces for two DNA hairpins and examined the time interval of the signals from the two fluorescent dyes (Table S5). Our experimental results revealed that most signals appeared and disappeared almost simultaneously (Figure 2B), with only 14.7% of the signals showing a time interval greater than one second (Figure 2C). This result underscores the high sensitivity of the ForceChrono probe.

**Figure 2.**
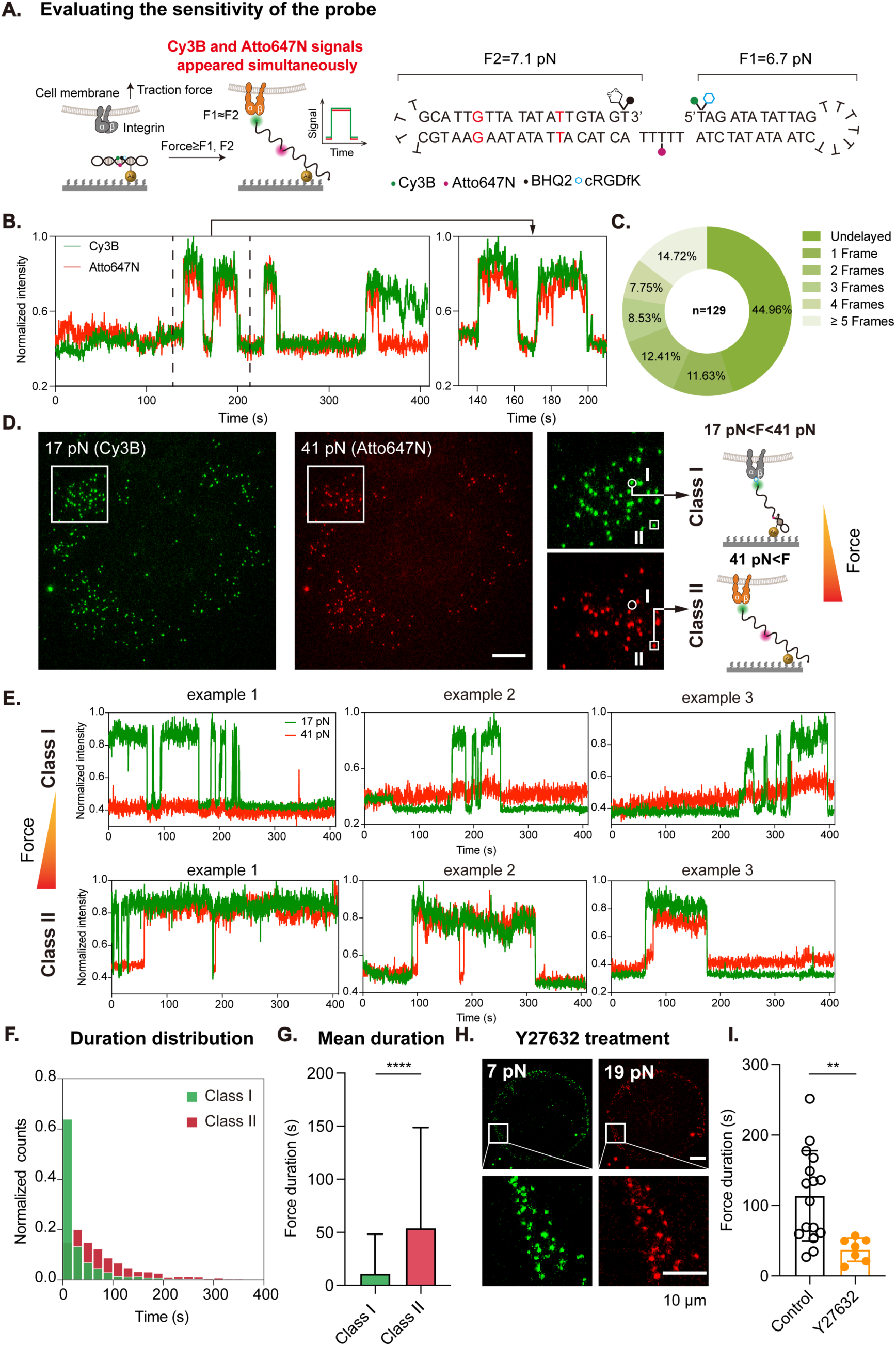
Investigating the relationship between magnitude and duration of integrin force. **A,** Evaluating the sensitivity of the probe. Left: The schematic illustrates that when the unfolding force of the two DNA hairpins is equal, the mechanical force on the integrin reaches the corresponding threshold, resulting in the simultaneous appearance of signals from the two fluorescent dyes. Right: The structure and oligonucleotide of the ForceChrono probe with the same unfolding force. **B,** A representative fluorescence signal trace of integrin force, showing that Cy3B and Atto647N signals appeared without delay. **C,** Measurement of delay frames between Cy3B and Atto647N signals. **D,** Left: Representative molecular force signal reported by 17-41 pN probes. Right: The schematic shows that integrins are divided into two classes according to the force magnitude. Class I integrins have a force between 17 pN and 41 pN, only Cy3B signals can be observed. Class II integrins have a force exceeding 41 pN, and Cy3B and Atto647N signals can be observed. **E,** Representative fluorescence intensity traces for these two integrins. **F,** Distribution of the duration of mechanical forces on these two integrins. **G,** Since the force duration of class II integrins cannot be fitted with an exponential function, the mean duration of a single integrin is shown here, based on data from over 1000 integrins. The error bars represent mean ± s.d. from three independent experiments. **H,** Representative 7-19 pN single-molecule fluorescence images after being treated with Y27632. **I,** Average duration of force on integrins with or without Y27632 treatment. Each point represents the mean duration of a cell. The error bars represent mean ± s.d. from three independent experiments. Two-tailed Student’s *t*-tests are used to assess statistical significance. Scale bar=10 μm.

Next, we used ForceChrono probes to classify integrins into two classes based on the magnitude of mechanical force they exerted (Figure 2D and S4A): Class I integrins exhibited a low magnitude of mechanical force (F1 < F < F2), while Class II integrins displayed high mechanical forces (F>F2). Interestingly, single-molecule fluorescence imaging revealed that Class I integrins are highly dynamic, transmitting lower magnitudes of force with shorter durations. In contrast, Class II integrins exhibited a different trend: the frequency of mechanical “blinking” events decreased, and the duration of force transmission increased. We observed intriguing behavior when employing the 17-41 pN ForceChrono probes (Figure 2E). At lower mechanical thresholds (17 pN < F < 41 pN), the mechanical force exerted on integrin-ligand bonds were predominantly unstable, manifesting short force durations. Intriguingly, upon exceeding a mechanical force of 41 pN, it exhibited remarkable stability. Upon analyzing the force duration distribution of the two integrin classes (Figure 2F), it was found that Class I integrins followed an exponential decay distribution. On the other hand, the distribution of Class II integrins decreased significantly in the 0-20 s duration range, while increasing in the long duration range. Further analysis of over 1000 integrins confirmed that Class II integrins had a longer average force duration compared to Class I integrins (Figure 2G). Similar results were obtained using 7-19 pN probes (Figure S4A-D). In contrast to the behavior of slip bonds, where bond stability decreases with increasing mechanical force ^34^, our findings more closely align with the characteristics of catch bonds. Our results reveal that the duration of force exerted on integrins increases with mechanical force up to a certain threshold.

To further validate our findings, we administered the ROCK inhibitor Y27632 to the cells, instigating a transition from Myosin II-driven forces to actin polymerization-driven forces (Figure 2H), which are intrinsically weaker. This pharmacological intervention effectively reduced the upper limit of mechanical forces exerted by individual integrins^10^, as evidenced by the diminished occurrence of forces exceeding 41 pN. We then used the 7-19 pN ForceChrono probe to investigate the dynamic characteristics of integrin force in Y27632-treated MEF cells, and we observed that the mechanical signals of Y27632-treated MEF cells were predominantly centered at cell edge locations, which is consistent with previous observations (Figure 2H)^10,35^. Compared to untreated cells, Y27632-treated cells exhibited a more dynamic mechano-fluorescence pattern with a significantly shorter overall force duration (Figure 2I), with each integrin force duration averaging about 20 seconds. This data further suggests that weaker mechanical forces make it difficult to stabilize the integrin-ligand bond.

Collectively, our data demonstrate the efficacy of ForceChrono probes in precisely quantifying single-molecule force durations in living cells. Our findings suggest a positive correlation between the magnitude and duration of mechanical force exerted by individual integrins. Importantly, we observed that lower mechanical forces are less effective in sustaining integrin-ligand mechanical transduction. This phenomenon is likely to contribute to the role of integrin as biomechanical sensors, facilitating the coupling of mechanical force with downstream biochemical signaling cascades, aligning with the catch-bond behavior previously observed in vitro studies^34^.

### The nonlinear dynamics of integrin loading rates

To comprehensively understand the dynamics of mechanical force loading on integrins, we employed two ForceChrono probes with distinct mechanical ranges: 7 pN-19 pN and 17 pN-41 pN. These probes offer a continuum of force thresholds, allowing for a nuanced investigation. Control experiments confirmed that the measured force durations were not significantly influenced by the probe structure, validating the comparability of the results across different probes (Figure S3E-G).

Our data revealed a striking heterogeneity in the loading rates of mechanical forces on integrins (Figure 3A and 3B). Specifically, after 2 hours of MEF cell spreading on surfaces modified with these probes, the loading rate was approximately 0.6 pN/s in the low-force range (7-19 pN). However, this rate escalated to about 1.5 pN/s in the 17-41 pN range. The nonlinear dynamic of the integrin loading rate was further confirmed in 3T3, C2C12, and A375 cells (Figure S4E-G), although the measured loading rates were not entirely consistent across different cell types.

**Figure 3.**
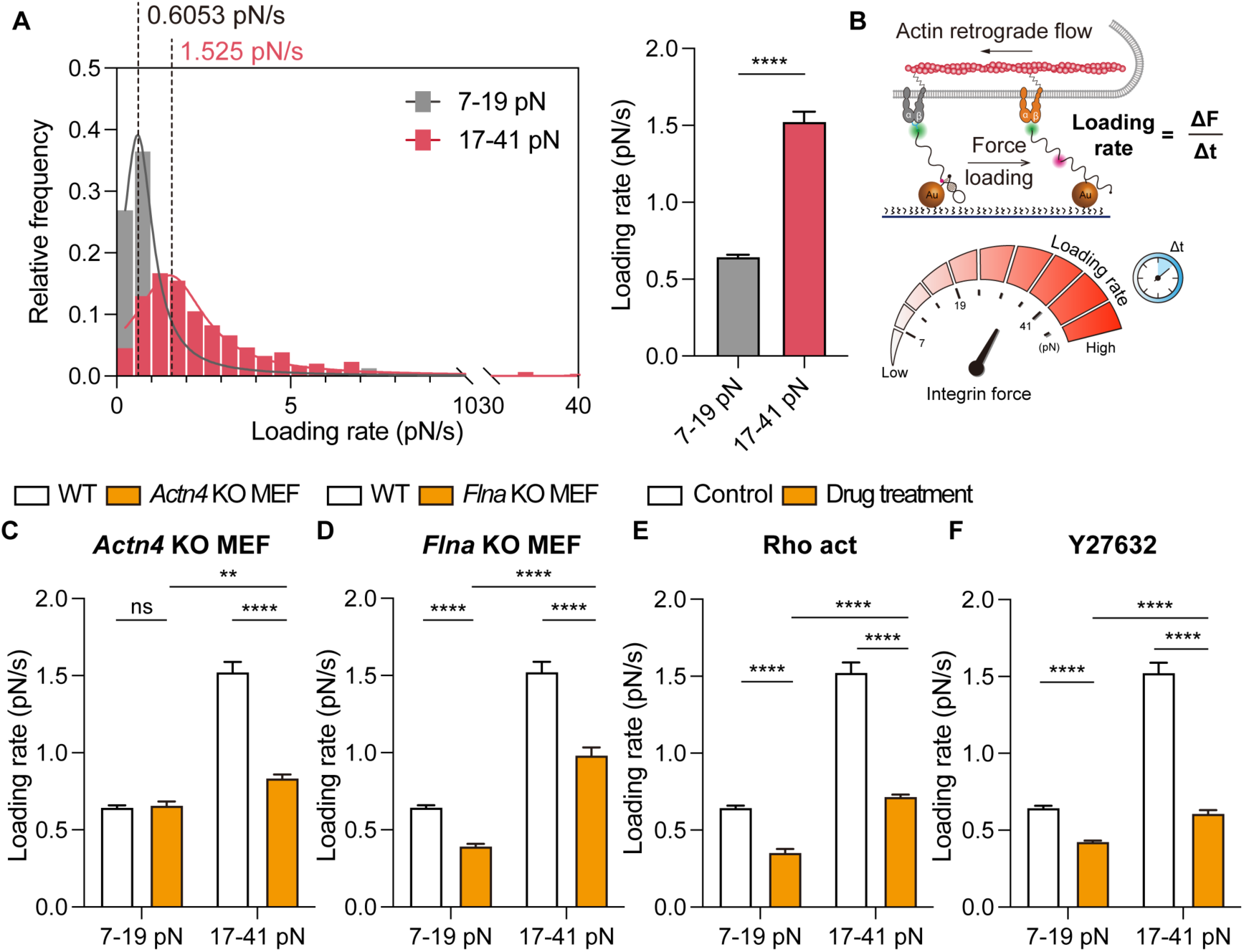
Integrin force loading rate in different mechanical magnitudes. **A,** Left: Distribution of integrin mechanical force loading rate in 7-19 pN and 17-41 pN ranges and the fitting results of Lorentzian distribution. Right: Average loading rate of force on integrins measured using 7-19 pN or 17-41 pN ForceChrono probes. The error bars represent mean ± 95% CI from three independent experiments. **B,** Upper: Diagram of measuring the loading rate of integrin force using ForceChrono probes. Lower: The schematic shows the relationship between the mechanical force magnitude of integrin and the loading rate. **C, D,** Loading rate of *Actn4* KO MEF (**C**) or *Flna* KO MEF (**D**) and WT MEF in different mechanical force magnitudes. **E, F,** Integrin force loading rate of MEF cells treated with Rho activator (**E**) or Y27632 (**F**). Two-tailed Student’s *t*-tests are used to assess statistical significance.

Given the crucial role of actin in the dynamics of integrin mechanical force loading, we investigated how filamin and α-actinin influence the mechanical loading rate of individual integrins. These two important actin-binding proteins are known for their ability to crosslink or bundle F-actin, forming specific actin structures: actin networks by filamin and actin bundles by α-actinin^36^. Additionally, they are involved in the linking and mechanical transduction of integrin and actin cytoskeleton^37,38^. To explore their impact, we knocked out filamin A or α-actinin-4 in MEF cells and studied their effects on integrin loading rates across various force intervals (Figure 3C and 3D). Our results showed that filamin A knockout decreased the loading rate across the 7-41 pN range, more noticeably in the lower force interval (7-19 pN). Conversely, α-actinin-4 knockout did not significantly affect the loading rate in the lower force interval but resulted in a decrease in the higher force interval (17-41 pN). This implies that filamin A contributes to integrin force loading across a broad range, whereas α-actinin-4’s influence becomes more pronounced at higher mechanical force levels. These findings suggest that the roles of filamin A and α-actinin-4 in integrin mechanical loading are distinct and depend on the force level, highlighting the nuanced interplay between cytoskeletal components and integrin mechanics.

Finally, we investigated the effect of myosin activity on the mechanical loading dynamics of integrin (Figure 3E and 3F). Our experiments, involving both Rho activator and Y27632 treatments, revealed that the integrin loading rate decreased significantly across both force ranges of 7-19 pN and 17-41 pN. This result suggests that myosin activity, a crucial factor in cytoskeletal dynamics and cellular force generation, plays a significant role in modulating integrin loading rates, impacting the overall mechanotransduction process.

Taken together, the compilation of our findings delineates a complex interplay between cellular architecture and integrin force transduction. One of the central to this relationship is the integrin force loading rate, a key parameter that bridges cellular mechano-sensitivity with the dynamics of integrin-mediated forces. This coupling dictates cellular mechano-sensitivity and significantly influences how cells respond and adapt within various mechanical environments, highlighting the intricate nature of cellular mechanotransduction processes.

### The vinculin-talin interaction affects the mechanical dynamics of integrin

Vinculin, a key player in the molecular clutch model, is pivotal for linking integrin-talin complexes to the actin cytoskeleton, facilitating dynamic engagement and disengagement with the extracellular substrate^39^. Despite extensive in vitro exploration of vinculin’s interaction with talin^20,23,24,40^, its specific impact on integrin-mediated force transmission within cells remains less understood.

We developed a vinculin-null mouse embryonic fibroblast (*Vcl* KO MEF) cell line^17^, stably expressing paxillin-GFP. Consistent with prior findings, vinculin absence did not impede focal adhesion formation or maturation (Figure 4A)^17,41^. However, our observations revealed that *Vcl* KO cells predominantly maintained an integrin force around 7 pN (Figure 4B), with a substantial decrease in integrins capable of exerting forces beyond 17 pN.

**Figure 4.**
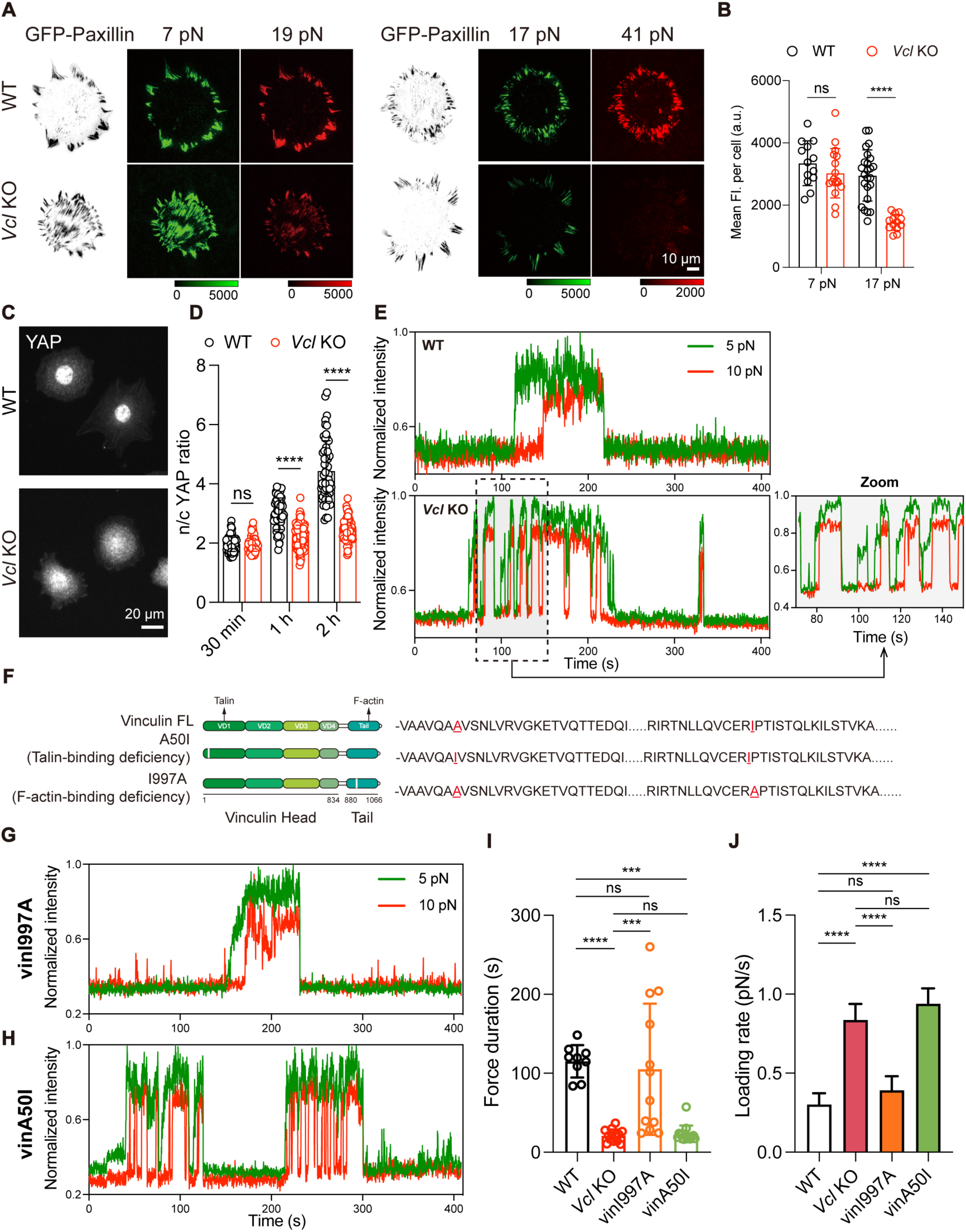
Role of vinculin in the mechanical transduction of integrin. **A,** Representative mechanical force images of GFP-paxillin stably expressed WT MEF or *Vcl* KO MEF cells on 7-19 pN or 17-41 pN ForceChrono probes. Scale bar=10 μm. **B,** Average intensity of tension signal in 7 pN or 17 pN channels for WT MEF or *Vcl* KO MEF cells. The error bars represent mean ± s.d. from three independent experiments. **C,** Representative images of YAP stainings of WT MEF or *Vcl* KO MEF cells spread out for 2 h. Scale bar=20 μm. **D,** n/c YAP ratio of cells spread out at different times. The error bars represent mean ± s.d. from three independent experiments. **E,** Representative fluorescence signal trace of WT MEF or *Vcl* KO MEF cells on 5-10 pN probes. **F,** Vinculin mutants: A50I (talin-binding deficient) and I997A (F-actin-binding deficient). **G, H,** Representative fluorescence signal trace of integrin force in vinI997A MEF (**G**) or vinA50I MEF (**H**). **I,** Average duration of force on integrins measured using 5-10 pN probes in WT, *Vcl* KO, vinI997A or vinA50I MEF cells. Each point represents the mean duration of a cell. The error bars represent mean ± s.d. from three independent experiments. **J**, Average loading rate of force on integrins measured using 5-10 pN probes in WT, *Vcl* KO, vinI997A or vinA50I MEF cells. The error bars represent mean ± 95% CI from three independent experiments. Two-tailed Student’s *t*-tests are used to assess statistical significance.

Additionally, we notice a significant difference in YAP nuclear translocation efficiency between *Vcl* KO and WT cells (Figure 4C and 4D). Previous studies have linked extracellular matrix (ECM)-mediated force transmission to nuclear flattening, facilitating nuclear pore stretching and lowering resistance to molecular transport such as YAP entry^42^. Since the vinculin is integral to connecting integrin-talin complexes to the actin cytoskeleton^43^, it is likely to play a crucial role in regulating the mechanical forces that facilitate YAP’s movement into the nucleus. Therefore, exploring the stability and duration of single integrin forces in the absence of vinculin informs our understanding of its mechanistic impact on cellular processes. Single-molecule ForceChrono probe imaging revealed stark contrasts in mechanical loading dynamics between WT and *Vcl* KO MEFs (Figure 4E, Movie S2 and S3). Vinculin-deficient cells showed transient, rapid fluctuations in single-molecule force signals, indicative of a higher loading rate (∼0.8 pN/s) and a shorter force duration (∼16 seconds). This suggests that vinculin absence disrupts the mechanical stability of the molecular clutch, leading to a “slipping” clutch mechanism with rapid force application and release. This mechanical instability in *Vcl* KO cells could explain the reduced YAP nuclear translocation efficiency, the transient and unstable mechanical forces might be insufficient for consistently stretching nuclear pores to the extent required for efficient YAP transport. Vinculin, therefore, emerges not just as a force transmitter but as a crucial regulator in the mechanotransduction pathway, influencing the nuclear translocation of mechanosensitive molecules like YAP by modulating the sustained application and stability of mechanical forces at the cellular level.

Further exploring vinculin’s domain-specific functions, we engineered two full-length vinculin mutants: vinA50I, deficient in talin-binding, and vinI997A, lacking actin-binding capabilities^43^, both GFP-tagged (Figure 4F). Reintroducing these mutants into *Vcl* KO MEFs, we observed that vinI997A partially restored integrin mechanical dynamics, approximating WT cell patterns (Figure 4G-J). This suggests that while the actin-binding domain is crucial^44^, it is not the exclusive contributor to integrin mechanical stability. In contrast, vinA50I exhibited mechanical dynamics akin to *Vcl* KO MEFs, underscoring the indispensable role of the talin-binding domain in maintaining force transmission and cellular adhesion. These insights deepen our understanding of vinculin’s domain-specific roles in integrin mechanotransduction and shed light on its integral function in cellular adhesion and mechanical equilibrium.

### Exploring the impact of ligand spacing on integrin mechanical dynamics

The density of ligands in the ECM plays a critical role in the formation of cellular adhesion structures, influencing cellular mechanosensing and behavior^45–47^. Previous research has established a critical threshold of 60-70 nm ligand spacing for the maturation of focal adhesions (FAs)^48,49^, which is essential for the force-mediated mechanosensing mechanism. However, the temporal dynamics of these forces, particularly how loading rate and duration vary with ligand spacing, have been less explored.

Using block copolymer micelle nanolithography (BCMN)^45,47^, we fabricated gold nanoparticle nanopatterned substrates with 40 nm and 100 nm ligand spacings to regulate the integrin ligand spacing (Figure S5). After functionalizing these substrates with 7-19 pN ForceChrono probes, we observed enhanced FA formation on the low-spacing (40 nm) substrate (Figure 5A), consistent with previous findings^35,47^.

**Figure 5.**
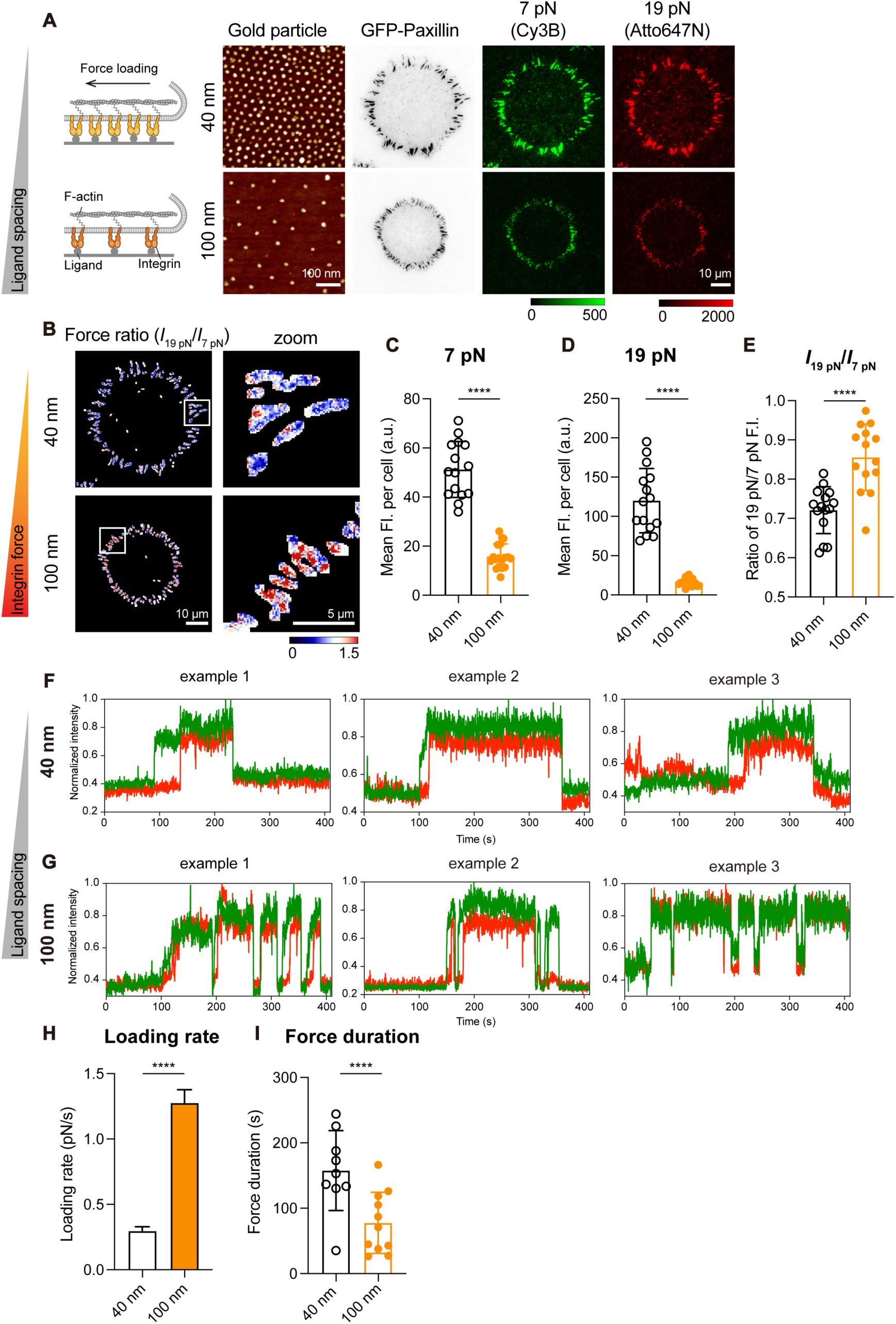
Ligand spacing affects the mechanical force of integrin. **A,** Representative AFM micrograph of the low or high-spacing gold particle ordered pattern substrates (Scale bar=100 nm), and the mechanical force signal reported by 7-19 pN probes after paxillin-GFP tagged MEF cells were plated onto these substrates (Scale bar=10 μm). **B,** Representative force ratio map (*I*19 pN/*I*7 pN) of cells at different ligand spacing substrates. Scale bar=10 μm (zoom, 5 μm). **C, D,** Average intensity of tension signal in 7 pN (Cy3B) or 19 pN (Atto647N) channels on these substrates. The error bars represent mean ± s.d. from three independent experiments. **E,** Fluorescence intensity ratio of the 19 pN to 7 pN. The error bars represent mean ± s.d. from three independent experiments. **F, G,** Representative fluorescence signal trace of integrin force in MEF cells at different ligand spacings. **H,** Average loading rate of force on integrins measured using 7-19 pN probes with different ligand spacings. The error bars represent mean ± 95% CI from three independent experiments. **I,** Average duration of force on integrins measured using 7-19 pN probes with different ligand spacings. The error bars represent mean ± s.d. from three independent experiments. Two-tailed Student’s *t*-tests are used to assess statistical significance.

The ForceChrono probe, featuring two series-connected probes with different threshold forces, allows accurate determination of integrin force ratios. The integrated structure of the ForceChrono probe ensures a more accurate and representative analysis of the mechanical forces involved. We, therefore, compared the difference in the mechanical ratio of cells on the substrates with different ligand spacing and found that on the high-spacing ligand patterned substrates (Figure 5B-E), the fluorescence intensity ratio of 19 pN to 7 pN was significantly higher than that of cells on the low-spacing ligand patterned substrates. This suggests that while increasing the ligand spacing affects the number of integrins activations and, ultimately, the focal adhesions, individual integrins increase the mechanical force at lower ligand densities. We then investigated the effect of ligand density on the mechanotransduction dynamics of individual integrins (Figure 5F-I). We found that at low ligand spacing, force fluorescence trajectories were stable, consistent with previous observations. Interestingly, compared to low ligand spacing, single-molecule signals at high ligand spacing exhibited transient and rapid flickering characteristics with an average duration of only 20 seconds, leading to shorter force duration and higher loading rates.

These findings suggest that ECM ligand density profoundly affects the magnitude and dynamics of integrin forces. At lower ligand densities, the forces exerted by myosin are distributed over fewer adhesions, increasing the load on each integrin. This not only accelerates force increase, but also facilitates focal adhesion disassembly, supporting recent predictions of molecular clutch models^45^.

## Discussion

In this study, we developed the DNA-based ForceChrono probe for analyzing integrin force dynamics at the single-molecule level in living cells. This probe not only quantifies the magnitude of single-molecule forces but also deciphers complex force application dynamics, including loading rate and duration.

The covalent bonding of the core components in the ForceChrono probe ensures structural precision and stability. Each ForceChrono probe molecule consists of two DNA hairpin probes that maintain their integrity under mechanical forces. This is in stark contrast to the annealing self-assembly method^50^, which is not suitable for single-molecule experiments due to structural instability and unreliable probe composition resulting from variations in annealing efficiency. Further, the ForceChrono probe’s method of calculating the loading rate by illuminating two distinct mechanical points as the monomolecular force increases effectively bypasses the problems associated with fluctuations in single-molecule fluorescence resonance energy transfer (smFRET) setups^51–53^, enhancing force measurement accuracy. Additionally, its dual-probe configuration enables more effective monitoring of force disappearance, reducing misinterpretation risks due to dye photobleaching. This approach captures molecular spring unfolding dynamics with improved accuracy and reliability.

Our findings significantly narrow the previously broad estimates of force loading rates exerted by living cells, which ranged from 0.007 to 10^7^ pN/s^27,30,31^, to a more precise range of approximately 0.5 to 2 pN/s. This finding brings clarity to the field of cellular mechanics, suggesting a typical range for cell membrane protein receptor-mediated force transduction. Moreover, the use of ForceChrono probes allows for the dissection of the role of various proteins in the mechanical dynamics of integrin, such as the crucial components of the molecular clutch. Notably, the study sheds light on the nonlinear nature of integrin loading rates, offering insights into the complex interplay of cellular components in integrin mechanodynamics and their role in mechanotransduction.

In addition, the experimental results from the ForceChrono probe have revealed a crucial relationship between the magnitude of integrin forces and their duration, highlighting a key aspect of cellular mechanotransduction. We observed that smaller integrin forces, approximately a few pN, correspond to shorter durations of integrin-ECM binding. Conversely, as these forces increase to higher magnitudes (e.g.,> 41 pN), integrin-ECM linkages exhibit greater stability. This finding is consistent with the behavior of catch bonds observed in in vitro studies^34^ and provides critical insights into the molecular dynamics of cellular adhesion and the mechanical stability of integrin-ECM interactions.

In summary, our results demonstrate that DNA-based ForceChrono probes provide multiple perspectives for studying the dynamics of mechanical loading in living cells. We believe that these probes will serve as powerful tools to elucidate the link between cellular mechanical sensing and downstream signaling cascades, including immune recognition, stem cell differentiation and tumor metastasis.

## Acknowledgments

We thank Professor Khalid Salaita for his feedback on the manuscript. This work was supported by the National Natural Science Foundation of China (21775115, 32150016, 32071305, 31871356), the Fundamental Research Funds for the Central Universities (2042021kf0030) and Innovation Funds for Postdocs in Hubei Province.

## Author contributions

Z.L. conceived the project and supervised the research. Y.H. and H.L. designed and conducted the overall experiments and analyzed the data with the help of others. C.Z. and X.Z. performed single-molecular magnetic tweezers experiments. J.F. provided the gold nanoparticle nanopatterned substrates and the *Actn4* KO MEF cell line. M.Y provided *Flna* KO MEF cell line. Z.L. and Y.H. wrote the paper.

## References

1. Shattil, S.J., Kim, C., and Ginsberg, M.H. (2010). The final steps of integrin activation: the end game. Nat. Rev. Mol. Cell Biol. 11, 288–300.

2. Iskratsch, T., Wolfenson, H., and Sheetz, M.P. (2014). Appreciating force and shape-the rise of mechanotransduction in cell biology. Nat. Rev. Mol. Cell Biol. 15, 825–833.

3. Hoffman, B.D., Grashoff, C., and Schwartz, M.A. (2011). Dynamic molecular processes mediate cellular mechanotransduction. Nature 475, 316–323.

4. Polacheck, W.J., and Chen, C.S. (2016). Measuring cell-generated forces: a guide to the available tools. Nat. Methods 13, 415–423.

5. Roca-Cusachs, P., Conte, V., and Trepat, X. (2017). Quantifying forces in cell biology. Nat. Cell Biol. 19, 742–751.

6. Stabley, D.R., Jurchenko, C., Marshall, S.S., and Salaita, K.S. (2011). Visualizing mechanical tension across membrane receptors with a fluorescent sensor. Nat. Methods 9, 64–67.

7. Liu, Y., Yehl, K., Narui, Y., and Salaita, K. (2013). Tension sensing nanoparticles for mechano-imaging at the living/nonliving interface. J. Am. Chem. Soc. 135, 5320–5323.

8. Zhang, Y., Ge, C., Zhu, C., and Salaita, K. (2014). DNA-based digital tension probes reveal integrin forces during early cell adhesion. Nat. Commun. 5, 5167.

9. Blakely, B.L., Dumelin, C.E., Trappmann, B., McGregor, L.M., Choi, C.K., Anthony, P.C., Duesterberg, V.K., Baker, B.M., Block, S.M., Liu, D.R., and Chen, C.S. (2014). A DNA-based molecular probe for optically reporting cellular traction forces. Nat. Methods 11, 1229–1232.

10. Li, H.Y., Zhang, C., Hu, Y.R., Liu, P.X., Sun, F., Chen, W., Zhang, X.H., Ma, J., Wang, W.X., Wang, L., et al. (2021). A reversible shearing DNA probe for visualizing mechanically strong receptors in living cells. Nat. Cell Biol. 23, 642–651.

11. Wang, X., and Ha, T. (2013). Defining single molecular forces required to activate integrin and notch signaling. Science 340, 991–994.

12. Duan, Y., Szlam, F., Hu, Y., Chen, W., Li, R., Ke, Y., Sniecinski, R., and Salaita, K. (2023). Detection of cellular traction forces via the force-triggered Cas12a-mediated catalytic cleavage of a fluorogenic reporter strand. Nat. Biomed. Eng. 7, 1404–1418.

13. Rao, T.C., Ma, V.P.Y., Blanchard, A., Urner, T.M., Grandhi, S., Salaita, K., and Mattheyses, A.L. (2020). EGFR activation attenuates the mechanical threshold for integrin tension and focal adhesion formation. J. Cell Sci. 133, jcs238840.

14. Hu, Y., Li, H., Wang, W., Sun, F., Wu, C., Chen, W., and Liu, Z. (2023). Molecular Force Imaging Reveals That Integrin-Dependent Mechanical Checkpoint Regulates Fcγ-Receptor-Mediated Phagocytosis in Macrophages. Nano Lett. 23, 5562–5572.

15. Liu, Y., Blanchfield, L., Ma, V.P.Y., Andargachew, R., Galior, K., Liu, Z., Evavold, B., and Salaita, K. (2016). DNA-based nanoparticle tension sensors reveal that T-cell receptors transmit defined pN forces to their antigens for enhanced fidelity. Proc. Natl. Acad. Sci. USA 113, 5610–5615.

16. Ma, V.P.Y., Hu, Y.S., Kellner, A.V., Brockman, J.M., Velusamy, A., Blanchard, A.T., Evavold, B.D., Alon, R., and Salaita, K. (2022). The magnitude of LFA-1/ICAM-1 forces fine-tune TCR-triggered T cell activation. Sci. Adv. 8, eabg4485.

17. Wang, W.X., Chen, W., Wu, C.Y., Zhang, C., Feng, J.J., Liu, P.X., Hu, Y.R., Li, H.Y., Sun, F., Jiang, K., et al. (2023). Hydrogel-based molecular tension fluorescence microscopy for investigating receptor-mediated rigidity sensing. Nat. Methods 20, 1780–1789.

18. Liu, B.Y., Chen, W., Evavold, B.D., and Zhu, C. (2014). Accumulation of dynamic catch bonds between TCR and agonist peptide-MHC triggers T Cell signaling. Cell 157, 357–368.

19. Marshall, B.T., Long, M., Piper, J.W., Yago, T., McEver, R.P., and Zhu, C. (2003). Direct observation of catch bonds involving cell-adhesion molecules. Nature 423, 190–193.

20. Yao, M., Goult, B.T., Klapholz, B., Hu, X., Toseland, C.P., Guo, Y., Cong, P., Sheetz, M.P., and Yan, J. (2016). The mechanical response of talin. Nat. Commun. 7, 11966.

21. Chen, W., Lou, J., Evans, E.A., and Zhu, C. (2012). Observing force-regulated conformational changes and ligand dissociation from a single integrin on cells. J. Cell Biol. 199, 497–512.

22. Chen, H., Yuan, G., Winardhi, R.S., Yao, M., Popa, I., Fernandez, J.M., and Yan, J. (2015). Dynamics of equilibrium folding and unfolding transitions of titin immunoglobulin domain under constant forces. J. Am. Chem. Soc. 137, 3540–3546.

23. Yao, M., Goult, B.T., Chen, H., Cong, P., Sheetz, M.P., and Yan, J. (2014). Mechanical activation of vinculin binding to talin locks talin in an unfolded conformation. Sci. Rep. 4, 4610.

24. Wang, Y.N., Yao, M.X., Baker, K.B., Gough, R.E., Le, S., Goult, B.T., and Yan, J. (2021). Force-Dependent Interactions between Talin and Full-Length Vinculin. J. Am. Chem. Soc. 143, 14726–14737.

25. Evans, E., and Ritchie, K. (1997). Dynamic strength of molecular adhesion bonds. Biophys. J. 72, 1541–1555.

26. Rief, M., Gautel, M., Oesterhelt, F., Fernandez, J.M., and Gaub, H.E. (1997). Reversible unfolding of individual titin immunoglobulin domains by AFM. Science 276, 1109–1112.

27. Jiang, L., Sun, Z., Chen, X., Li, J., Xu, Y., Zu, Y., Hu, J., Han, D., and Yang, C. (2016). Cells Sensing Mechanical Cues: Stiffness Influences the Lifetime of Cell-Extracellular Matrix Interactions by Affecting the Loading Rate. ACS Nano 10, 207–217.

28. Andreu, I., Falcones, B., Hurst, S., Chahare, N., Quiroga, X., Le Roux, A.L., Kechagia, Z., Beedle, A.E.M., Elosegui-Artola, A., Trepat, X., et al. (2021). The force loading rate drives cell mechanosensing through both reinforcement and cytoskeletal softening. Nat. Commun. 12, 4229.

29. Merkel, R., Nassoy, P., Leung, A., Ritchie, K., and Evans, E. (1999). Energy landscapes of receptor-ligand bonds explored with dynamic force spectroscopy. Nature 397, 50–53.

30. Moore, S.W., Roca-Cusachs, P., and Sheetz, M.P. (2010). S Stretchy proteins on stretchy substrates: the important elements of integrin-mediated rigidity sensing. Dev. Cell 19, 194–206.

31. Amar, K., Suni, II, and Chowdhury, F. (2020). A quartz crystal microbalance based study reveals living cell loading rate via αvβ3 integrins. Biochem. Biophys. Res. Commun. 524, 1051–1056.

32. Yun, C.S., Javier, A., Jennings, T., Fisher, M., Hira, S., Peterson, S., Hopkins, B., Reich, N.O., and Strouse, G.F. (2005). Nanometal surface energy transfer in optical rulers, breaking the FRET barrier. J. Am. Chem. Soc. 127, 3115–3119.

33. Bodescu, M.A., Aretz, J., Grison, M., Rief, M., and Fassler, R. (2023). Kindlin stabilizes the talin.integrin bond under mechanical load by generating an ideal bond. Proc. Natl. Acad. Sci. USA 120, e2218116120.

34. Kong, F., García, A.J., Mould, A.P., Humphries, M.J., and Zhu, C. (2009). Demonstration of catch bonds between an integrin and its ligand. J. Cell Biol. 185, 1275–1284.

35. Liu, Y., Medda, R., Liu, Z., Galior, K., Yehl, K., Spatz, J.P., Cavalcanti-Adam, E.A., and Salaita, K. (2014). Nanoparticle tension probes patterned at the nanoscale: impact of integrin clustering on force transmission. Nano Lett. 14, 5539–5546.

36. Ferrer, J.M., Lee, H., Chen, J., Pelz, B., Nakamura, F., Kamm, R.D., and Lang, M.J. (2008). Measuring molecular rupture forces between single actin filaments and actin-binding proteins. Proc. Natl. Acad. Sci. USA 105, 9221–9226.

37. Roca-Cusachs, P., del Rio, A., Puklin-Faucher, E., Gauthier, N.C., Biais, N., and Sheetz, M.P. (2013). Integrin-dependent force transmission to the extracellular matrix by alpha-actinin triggers adhesion maturation. Proc. Natl. Acad. Sci. USA 110, E1361–1370.

38. Kiema, T., Lad, Y., Jiang, P., Oxley, C.L., Baldassarre, M., Wegener, K.L., Campbell, I.D., Ylanne, J., and Calderwood, D.A. (2006). The molecular basis of filamin binding to integrins and competition with talin. Mol. Cell 21, 337–347.

39. Elosegui-Artola, A., Trepat, X., and Roca-Cusachs, P. (2018). Control of Mechanotransduction by Molecular Clutch Dynamics. Trends Cell Biol. 28, 356–367.

40. del Rio, A., Perez-Jimenez, R., Liu, R., Roca-Cusachs, P., Fernandez, J.M., and Sheetz, M.P. (2009). Stretching single talin rod molecules activates vinculin binding. Science 323, 638–641.

41. Austin, J., Tu, Y., Pal, K., and Wang, X.F. (2023). Vinculin transmits high-level integrin tensions that are dispensable for focal adhesion formation. Biophys. J. 122, 156–167.

42. Elosegui-Artola, A., Andreu, I., Beedle, A.E.M., Lezamiz, A., Uroz, M., Kosmalska, A.J., Oria, R., Kechagia, J.Z., Rico-Lastres, P., Le Roux, A.L., et al. (2017). Force Triggers YAP Nuclear Entry by Regulating Transport across Nuclear Pores. Cell 171, 1397–1410 e1314.

43. Atherton, P., Stutchbury, B., Wang, D.Y., Jethwa, D., Tsang, R., Meiler-Rodriguez, E., Wang, P., Bate, N., Zent, R., Barsukov, I.L., et al. (2015). Vinculin controls talin engagement with the actomyosin machinery. Nat. Commun. 6, 10038.

44. Thievessen, I., Thompson, P.M., Berlemont, S., Plevock, K.M., Plotnikov, S.V., Zemljic-Harpf, A., Ross, R.S., Davidson, M.W., Danuser, G., Campbell, S.L., et al. (2013). Vinculin-actin interaction couples actin retrograde flow to focal adhesions, but is dispensable for focal adhesion growth. J. Cell Biol. 202, 163–177.

45. Oria, R., Wiegand, T., Escribano, J., Elosegui-Artola, A., Uriarte, J.J., Moreno-Pulido, C., Platzman, I., Delcanale, P., Albertazzi, L., Navajas, D., et al. (2017). Force loading explains spatial sensing of ligands by cells. Nature 552, 219–224.

46. Cavalcanti-Adam, E.A., Micoulet, A., Blummel, J., Auernheimer, J., Kessler, H., and Spatz, J.P. (2006). Lateral spacing of integrin ligands influences cell spreading and focal adhesion assembly. Eur. J. Cell Biol. 85, 219–224.

47. Cavalcanti-Adam, E.A., Volberg, T., Micoulet, A., Kessler, H., Geiger, B., and Spatz, J.P. (2007). Cell spreading and focal adhesion dynamics are regulated by spacing of integrin ligands. Biophys. J. 92, 2964–2974.

48. Schvartzman, M., Palma, M., Sable, J., Abramson, J., Hu, X., Sheetz, M.P., and Wind, S.J. (2011). Nanolithographic control of the spatial organization of cellular adhesion receptors at the single-molecule level. Nano Lett. 11, 1306–1312.

49. Huang, J., Grater, S.V., Corbellini, F., Rinck, S., Bock, E., Kemkemer, R., Kessler, H., Ding, J., and Spatz, J.P. (2009). Impact of order and disorder in RGD nanopatterns on cell adhesion. Nano Lett. 9, 1111–1116.

50. Zhao, B., Li, N.W., Xie, T.F., Bagheri, Y., Liang, C.W., Keshri, P., Sun, Y.B., and You, M.X. (2020). Quantifying tensile forces at cell-cell junctions with a DNA-based fluorescent probe. Chem. Sci. 11, 8558–8566.

51. Göhring, J., Kellner, F., Schrangl, L., Platzer, R., Klotzsch, E., Stockinger, H., Huppa, J.B., and Schütz, G.J. (2021). Temporal analysis of T-cell receptor-imposed forces via quantitative single molecule FRET measurements. Nat. Commun. 12, 2502.

52. Tan, S.J., Chang, A.C., Miller, C.M., Anderson, S.M., Prahl, L.S., Odde, D.J., and Dunn, A.R. (2020). Regulation and dynamics of force transmission at individual cell-matrix adhesion bonds. Sci. Adv. 6, eaax0317.

53. Morimatsu, M., Mekhdjian, A.H., Adhikari, A.S., and Dunn, A.R. (2013). Molecular tension sensors report forces generated by single integrin molecules in living cells. Nano Lett. 13, 3985–3989.

