## Supporting Information for "DNA-based ForceChrono Probes for Deciphering Single-Molecule Force Dynamics in Living Cells"

### Table of contents:

**Methods**
**Table S1:** Sequence of probes (5'-3').
**Table S2:** Calculated the unzipping force ($F_{1/2}$) of the hairpin during different conditions.
**Table S3:** Employed various models to calculate the shearing force required for rupture dsDNA.
**Table S4:** Summary of the thresholds of ForceChrono probes.
**Table S5:** Calculated the unzipping force ($F_{1/2}$) of the second hairpin of Bi-7 pN probe.
**Figure S1:** Structure and synthesis scheme of ForceChrono probes.
**Figure S2:** Calibration of ForceChrono probes using magnetic tweezers.
**Figure S3:** Evaluating the stability and sensitivity of ForceChrono probes.
**Figure S4:** Investigation of the mechanical force duration and loading rate on integrins using ForceChrono probes.
**Figure S5:** Characterization of nanopatterned AuNP arrays.
**Movie captions**

### Methods

#### Cell culture and transfection

MEF (SCSP-105), C2C12 (SCSP-505), NIH-3T3 (SCSP-515) and A375 (SCSP-533) cells were purchased from Stem Cell Bank, Chinese Academy of Science. MEF, C2C12 and A375 cells were cultured in Dulbecco's Modified Eagle's Medium (DMEM, Sangon Biotech, E600003) supplemented with 10% fetal bovine serum (FBS, VivaCell, C04001), 100 U $mL^{-1}$ penicillin-streptomycin (Sangon Biotech, E607011) and incubated at 37 °C and 5% $CO_2$. NIH-3T3 cells were cultured in DMEM supplemented with 10% newborn calf serum (CS, Gibco, 16010159), 100 U $mL^{-1}$ penicillin-streptomycin, and incubated at 37 °C and 5% $CO_2$.

Vinculin knockout MEF cells were generated and cultured as previously described[1]. To generate $\alpha$-actinin-4 or filamin A knockout MEF cells using CRISPR, the single guide RNA sequences were designed to target *Actn4* or *Flna* gene and cloned into pSpCas9(BB)-2A-Puro (PX459) (Addgene, 48139). According to the manufacturer's protocol, all transfections were carried out using the Lipofectamine™ 3000 Transfection Reagent (Thermo Fisher, L3000075).

sgRNA1 for *Actn4* gene: 5'-CGGAAGCATGGGCGACTACA-3'

sgRNA2 for *Actn4* gene: 5'-AGCGTTTGCCTAAGCCAGAG-3'

sgRNA1 for *Flna* gene: 5'-GCAGCTCGAAAATGTGTCGG-3'

sgRNA2 for *Flna* gene: 5'-CAGTATCGATTCACGGGACG-3'

#### DNA constructs and antibodies

The following antibodies were used: Anti-alpha Actinin-4 (Abcam, ab108198); Anti-Filamin A (Abcam, ab76289); Anti-GAPDH mouse monoclonal antibody (Sangon Biotech, D190090); HRP-conjugated Goat Anti-Rabbit IgG (Sangon Biotech, D110058); HRP-conjugated Goat Anti-Mouse IgG (Sangon Biotech, D110087).

eGFP-LifeAct and eGFP-Paxillin Lentiviral expression plasmids were gifts from

Prof. Congying Wu's lab (Peking University, China). pSpCas9(BB)-2A-Puro (PX459) plasmid was a gift from Prof. Kai Jiang's lab (Wuhan University, China). For the expression in vinculin knockout MEF cells, the vinculin mutant vinA50I-eGFP and vinI997A-eGFP were generated by polymerase chain reaction method and cloned into PLVX plasmids using the Seamless Cloning Kit (Beyotime, D7010M) according to the manufacturer's protocol, and verified by DNA sequencing.

**Western Blots**

Western blots were performed to verify the knockout efficiency of proteins. In Brief, cells were washed with PBS and lysed in RIPA buffer (Sangon Biotech, C500005) with Protease inhibitor (Roche, 04693159001). Protein concentration was measured using the BCA Protein Assay Kit (Sangon Biotech, C503061) and normalized across samples. After being mixed with loading buffer and boiled at 95°C for 5 min, protein lysates were separated on 4%-20% gradient SDS-PAGE gels and transferred to PVDF membranes (Millipore, IPVH00010). Membranes were blocked with 5% w/v nonfat milk in TBST, and immunoblot analysis was performed with the corresponding primary antibodies and HRP-conjugated secondary antibodies. Protein bands were detected with ECL reagents (Beyotime, P0018AS) and visualized using the Bio-Rad Gel Doc XR+ System.

**Fluorescence immunostaining**

For immunostaining of YAP in WT MEFs and Vcl KO MEFs, cells were fixed with 4% PFA (Polyformaldehyde) at 37°C after spreading out for different time, permeabilized for 10 minutes with 0.2% Triton X-100 (Sigma-Aldrich) at room temperature, after a 1 hour blocking step with 3% BSA (Bovine Serum Albumin), a 1:200 dilution of mouse monoclonal anti-YAP1 (Santa Cruz Biotechnology, sc-101199) was used as primary antibody and incubated with cells at 4°C overnight. The cells were then incubated with TRITC-labeled anti-mouse secondary antibody (Sangon Biotech, D110083, 1:200 dilution) and FITClabeled phalloidin (Solarbio, CA1620, 1:300 dilution) at room temperature for 1 hour, respectively. Finally, the nuclei were labeled with DAPI (Sangon Biotech, E607303), and all stained specimens were rinsed extensively with PBS before microscopic observations.

**Synthesis of ForeChrono probes**

The ForceChrono probes were assembled from three DNA strands: strand A, strand B, and strand C (Table S1). Briefly, they were custom-synthesized and purified by Sangon Biotech. Strand A was modified with CH-CH groups and C6-$NH_2$ at its 5'-end. Strand B was modified with a phosphate group and C6-$NH_2$ at its 5'-end and internal position, respectively. Strand C was modified with a phosphate group at its 5'-end, and C6-$NH_2$ and BHQ2 quencher were modified at its 3'-end. The step-by-step synthesis procedure is as follows:

Step 1 (Synthesis of strand A). First, strand A was coupled with a fluorescent dye (Cy3B) via the amide linkage formed between the N-Hydroxysuccinimide (NHS) ester group on the dye and the modified amine group on the DNA strand. 20 µL DNA (0.2 mM) in $H_2O$ was mixed with 2.5 µL Cy3B-NHS ester (10 mM) diluted in DMSO, then added 2.5 µL $NaHCO_3$ (1 M) to adjust pH to 8.3. The mixture was incubated overnight at room temperature. After a rough separation through a P6 gel desalting column (Bio-Rad, 7326002) to remove the residual dye, the dye-labeled DNA was further separated by HPLC (elution A: 0.1 M TEAA, pH 7.0, elution B: acetonitrile; gradient:10-60% elution B over 50 min; flow rate:1 mL/min). After dye labeling, the next step was to couple a cyclic RGDfK peptide to the DNA-dye conjugate by a copper-catalyzed "click" reaction between the azide on the peptide and the alkyne on the DNA. Finally, the dye-DNA-peptide conjugate was purified by 15% denaturing urea polyacrylamide gel electrophoresis. The desired gel band was cut and crushed, then the crushed gel was soaked in PBS and vortexed for 4 h at room temperature. After filtered through a 0.22 µm centrifugal filter (Corning, 8162-ZX) to remove gels, the product was purified through a P-6 gel desalting column.

Step 2 (Synthesis of strand B). Strand B was labeled with an Atto647N dye by coupling the amine group on the DNA and the NHS ester on the Atto647N. The reaction and purification of Atto647N-labeled stand B were performed as described in step 1.

Step 3 (Synthesis of strand C). Strand C was reacted with NHS ester terminal lipoic acid (CAS, 40846-94-4) and purified through HPLC, as described in step 1.

Step 4 (Achieving the final probes). The products obtained above were mixed in PBS at a molar ratio of 1:1.1:1 (strand A: strand B: strand C), heated at 95℃ for 5 min, and cooled to room temperature. After the segments were annealed, T4 DNA ligase (Sangon Biotech, B522241) was used to seal the nicks between the segments following the manufacturer's protocols. The final products were separated and purified by 10% denaturing urea polyacrylamide gel electrophoresis. Before use, another anneal step was needed to make the oligonucleotide form the correct secondary structure.

**Estimation of rupture force of ForceChrono probes**

Because magnetic tweezers and cell force imaging were performed at different conditions, 22°C , PBS and 37°C, DMEM cell culture medium, respectively. The forces to unfold the probe under different temperatures and ionic concentrations would differ. We inferred the unfolding force of the probe according to previous publications.

The unzipping force ($F_{1/2}$) of the hairpin domain of the probe at different conditions (**a**, 22°C, PBS: 157 mM $Na^+$; **b**, 37°C, DMEM culture: 155.3 mM $Na^+$, 0.8 mM $Mg^{2+}$) was calculated as described by Woodside and colleagues[2]:

$$F_{1/2} = \frac{\Delta G_{fold} + \Delta G_{stretch}}{\Delta_x}$$

Where,

$$\Delta G_{stretch} = \frac{k_B T}{L_p} \frac{L_c}{4(1 - x/L_c)} [3\left(\frac{x}{L_c}\right)^2 - 2\left(\frac{x}{L_c}\right)^3]$$

$\Delta_x$ is the displacement of the two ends of the hairpin under the force of $F_{1/2}$ ($\Delta_x$=(0.44n-2) nm; n, length of the hairpin in nt). $k_B$ is the Boltzmann constant. $T$ is the temperature. $x$ is the ssDNA extension at $F_{1/2}$ (0.44 nm/nt). $L_c$ is the contour length of ssDNA (0.63 nm/nt).

$\Delta G_{fold}$ was calculated using the IDT oligoanalyzer 3.1 and was shown in Table S2.

When the force is applied to the hairpin stem in a shear geometry, the shearing force to rupture the duplex will increase with the duplex length within a specific range. Several models have been used to explain the shear force required to rupture the double-stranded DNA. We found that the model proposed by de Gennes and the modified form by K. Hatch can be well fit to the system in which the force is loaded dynamically[3,4], despite that this model ignores the effect of the duration of force exertion. Based on the de Gennes model, the critical force to rupture a DNA duplex is,

$$f_c = 2f_1\left[\chi^{-1}\tanh\left(\frac{\chi N}{2}\right) + 1\right]$$

K. Hatch introduced a parameter $N_{open}$ (7 bp) to account for the effect of thermal energy, which keeps several base pairs at the end in the open state, thereby constraining the actual double-stand length to N-$N_{open}$[4]. So, the modified equation is,

$$f_c = 2f_1\left[\chi^{-1}\tanh\left(\frac{\chi(N - N_{open})}{2}\right) + 1\right]$$

In the mixed model where the force at one end was exerted on the internal nucleotide rather than on the end, $N_{open}$ should be halved, resulting in 3.5 bp. Another important factor in determining the critical force value to rupture a DNA duplex is the observation time, indicating that even small forces can break an extended DNA duplex given sufficient time. In our previous work, we utilized the toy model described by Majid Mosayebi et al. to estimate the shear force needed to rupture the DNA duplex in the hairpin stem[5].

$$f_c = \frac{N\Delta G_{bp} - \Delta G_{\tau_{obs}} - \Delta G_0}{N\delta - \delta_0}$$

The parameters used in the publication for the rupture of a DNA duplex by applying antiparallel force on the 3'-3' or 5'-5' ends were based on the measurement by K. Hatch[4]. Where $N$ is the number of base pairs under shearing, $\Delta G_{bp}$ is the free-energy cost to break the base pair (1.5 kcal/mol), $\delta$ is increased extension per base pair of the DNA duplex in the transition state relative to the fully formed duplex (0.14 nm per bp). The transition state was assumed to be a single base-pair state, so $\delta_0$, the offset that represents the base pairs in the transition state, is equal to $\delta$, and $\Delta G_0$ is equal to $\Delta G_{bp}$. $\Delta G_{\tau_{obs}}$ is the free-energy barrier that needs to be surmounted between the fully formed duplex and the transition state (10.4 kcal/mol) during the observation time (1 s). $\Delta G_{\tau_{obs}}$ introduced the influence of temperature and observation time, indicating that under the critical force, observation time and temperature, it results in a 50% rupture probability.

In the mixed model, where the force at one end was exerted on the internal nucleotide rather than on the end. The presence of the loop and residual duplex contributes to increased stability in the stem under force stretching. As a result, we propose that a higher force may be required to lower the energy barrier $\Delta G_{\tau_{obs}}$. We speculate that a hairpin with a 3-4 bp stem will open spontaneously at room temperature, much shorter than the DNA duplex. Therefore, the $\Delta G_{\tau_{obs}}$ will be 3-4 $\Delta G_{bp}$. The unfolding force of the hairpin structure calculated according to different models is shown in Table S3.

**Magnetic tweezers calibrate ForceChrono probes**

The DNA construct for the magnetic tweezers included a force probe domain tethered to the glass substrate via a long double-strand DNA handle (4.7 kb). In brief, multi-digoxigenin labeled oligonucleotide was made from single primer PCR and was used as a primer to synthesize the long DNA handle by PCR amplification from a λDNA. Subsequently, the DNA handle was digested with Bsa I, leaving a 5' overhang with a CGGC sequence. DNA probes, containing a biotin at the same site as cRGDfK, a GCCG overhang, and a phosphate group at its 5' end, were ligated to the DNA handle using T4 ligase. The entire DNA structure was attached to the anti-digoxigenin-coated glass substrate and the streptavidin-coated magnetic beads, respectively. The performance of single-molecule magnetic tweezers calibration experiments was basically consistent with the previously reported protocol[6]. Unless otherwise specified, all measurements were carried out at 22 °C in PBS, and the permanent magnets exerted a pulling force on the magnetic beads at a loading rate of 1 pN/s (Table S5). Additionally, the experiment was also carried out at different temperatures (30°C and 35°C) or changed the loading rate of force applied to the magnetic beads (from 0.1 pN/s to 4 pN/s) to calibrate the probes.

**Synthesis of gold nanoparticles**

The synthesis of 3.5 nm citrate-stabilized gold nanoparticles (AuNPs) was primarily based on the method described by Puntes et al.[7]. Briefly, 1 mL of $HAuCl_4$ (25 mM) was injected into a reaction solution containing 150 mL of sodium citrate (2.2 mM), 0.1 mL of tannic acid (2.5 mM) and 1 mL of potassium carbonate ($K_2CO_3$, 150 mM) at 70ºC. The reaction was completed in 5 min, and the extinction spectrum of AuNP was measured using a NanoDrop microvolume spectrophotometer (Thermo Fisher).

**Surface preparation**

The procedure for glass substrate functionalization was modified from previously published protocols[6,8]. Briefly, circular coverslips (25 mm diameter, 170 μm thickness) were first rinsed and sonicated three times in nanopure water (18.2 MΩ·cm) and then sonicated etched in 1 M KOH for 30 min. The temperature of the KOH should not exceed 40°C during the etching process. The coverslips were subsequently cleaned several times with nanopure water and sonicated three times in ethanol. After drying in an oven, the coverslips were hydroxylated in the oxygen plasma for 450 s and then incubated in ethanol with a 0.1% v/v (3-Aminopropyl) triethoxysilane for 1 h. The coverslips were alternately rinsed with ethanol and acetone, then dried under a stream of $N_2$. Subsequently, the coverslips were annealed for 1 h at 80°C. They were then reacted with 0.5% (w/v) lipoic acid-PEG-NHS (MW 3400) and 5% (w/v) mPEG-NHS (MW 2000) in 0.5 M $K_2SO_4$ and 0.1 M $NaHCO_3$ (pH=9) overnight at 4°C. After washing the coverslips with nanopure water, they were further passivated with T20 solution (5% v/v Tween-20 in 10 mM Tris-HCl with 50 mM NaCl, pH=8) for 30 min. Next, coverslips were washed with nanopure water and incubated with 150 nM 3.5 nm AuNP for 30 min. After washing the unreacted AuNP, the coverslips were dried under a steam of $N_2$ and placed in a nitrogen-filled dish, where they can be stored at 4°C for up to a week before use.

Lipoic acid-modified cRGDfK was mixed with ForceChrono probes in different proportions, and ForceChrono probes were gradient diluted to achieve single-molecule concentration. The mixture was added to the AuNP immobilization surfaces and incubated at room temperature for 30 min. After washing the coverslips with PBS, Four-color FluoSpheres microbeads were diluted with PBS at a ratio of 1:2500 and incubated for 10 min, serving as fiducial markers during data processing. These modified coverslips were then assembled into cell imaging chambers and immediately used for cell experiments.

**Microscopy and single-molecule fluorescence imaging for ForceChrono probes**

All images were obtained using a Nikon Eclipse Ti2 inverted microscope equipped with Nikon LU-N4 laser units featuring four lasers (405 nm, 488 nm, 561 nm, and 640 nm) with a power output of 15 mW. The microscope is also equipped with an autofocus system, a×100 oil objective (Nikon, 1.49 NA) and two ANDOR EMCCD (1024×1024 pixels) cameras. A dichroic short-wavelength pass filter (EO Edmund, 69217) was added to the optical path, reflecting 675-850 nm and transmitting 400-630 nm, enabling the two EMCCDs to simultaneously capture emissions from Cy3B and Atto647N, respectively. A stage top chamber (Okolab) was used, fitting into the x-y stage of the microscope, connecting to Okolab temperature, gas, and humidity controllers, and creating the appropriate environment (37°C, 5% $CO_2$) for live-cell imaging on the microscope stage. An objective warming apparatus was used to minimize the focus drift caused by the temperature changes. ForceChrono probes-modified surfaces were placed into a cell imaging chamber and prepared for cell spreading.

Images were acquired using Nikon NOS-Elements software with RAM capture. Simultaneously excitation of Cy3B and Atto647N was achieved using 561 nm and 640 nm lasers with powers of 1.5 mW and 4.5 mW, respectively. The two cameras were operated with an electron-multiplying gain of 300, and an exposure time of 200 ms was set, with a continuous capture of 2000 frames for each channel. Excitation of eGFP was accomplished using a 488 nm laser with a power of 1.5 mW and an exposure time of 200 ms.

When imaging, use the phenol red-free complete culture medium containing oxygen scavenging system (1 mg/mL of catalase and glucose oxidase), as well as 10 mM Trolox.

**Image processing**

Analysis of single-molecule fluorescent-time trajectories was performed using MATLAB 2019a and Image J. The process involved a three-step procedure: (1) Calibration using four-color microbeads to account for the deviation in two fluorescence channels caused by the two EMCCD images. The microbeads also helped to calibrate the x-y drift. (2) Selection of the region to be analyzed and subtraction of background noise. (3) Localization of the single-molecule fluorescent spots and extraction of the fluorescent-time trajectories.

**Force duration and loading rate data acquired and analysis**

The force duration measured by the ForceChrono probe was calculated by multiplying the number of frames of the Cy3B fluorescent signal by the time interval between each frame. As described above, 200 ms was used as the exposure time per frame in the experiment operation, but due to the delay, the EMCCD acquired each frame at an interval of 204 ms. It is worth noting that the time the probe was in the unfolded state can represent the force transmission duration on the integrin-ligand bond. Considering the cost of time from the initial formation of the bond to the unfolding of the first hairpin structure, the probe unfolded state represented the minimal bond lifetime. The duration distribution of mechanical force exerted on integrin was fitting with an exponential function:

$$y = y_0 + A_0 e^{-\frac{t}{\tau}}, y_0 > 0$$

Where $\tau$ is the mean force duration.

The loading rate of a single integrin mechanical force can be calculated by the time difference between opening the two hairpins of the probe after knowing the threshold of opening them. The Experimental data of loading rate was fitted by nonlinear curve-fitting in the least-squares using Lorentzian distribution and obtained the mean value of loading rate.

**Preparation of nanopatterned surface**

In order to regulate the integrin ligand density, we employed the nanopattern surface preparation method as previously described using block copolymer micelle nanolithography (BCMN)[9,10]. Specifically, polystyrene(x)-b-poly(2vinylpyridine)(y) diblock copolymers (PS-b-P2VP, PolymerSource Inc.) were dissolved with anhydrous toluene, and stirred at room temperature at 450 rpm for 24 h. Subsequently, $HAuCl_4 \cdot 3H_2O$ (Alfa Aesar) was added to the micellar solutions with a specific loading parameter defined as L = n[$HAuCL_4$]/n[P2VP] and stirred for an additional 24 h. Glass slides were pre-treated with piranha solution (98% $H_2SO_4$: 30% $H_2O_2$ = 3:1, v/v) and thoroughly rinsed with Milli-Q water, and then dipped into the micelle solution and pulled out to form a closely packed monolayer of micelles on the glass surfaces. These surfaces were then treated with oxygen plasma for 30 min to remove the polymer matrix, followed by immersion in ethanol for 30 minutes to facilitate reduction. After drying at 80 °C, the coverslips were stored for future use.

Before being used to image, the nanopatterned surfaces were cleaned by alternating washes with ethanol. After drying, the probe modification and micromanipulation procedures remained consistent with the substrate preparation methods. Furthermore, the PLL-g-PEG was incubated at room temperature for 30 min to block non-specific binding sites.

The surface morphology of the nano-gold pattern was characterized by AFM (BioScope Resolve, Bruker) and a silicon nitride AFM probe (SCANASYST-AIR, Bruker). The spacing of the gold particles was calculated by the mean nearest neighbor distance between five gold particles.

#### Statistical analysis

Data from experiments were graphed using GraphPad Prism 9 or MATLAB 2019a. Each point in the force duration statistical results represents the mean value fitted from a cell, and the statistical results of different conditions were presented in the form of mean ± s.d. unless otherwise noted. The loading rate statistics represent the mean value of the loading rate of all experimental data from individual integrins fitted by the Lorentzian distribution and the 95% confidence interval of the mean value. The Student's *t*-test was performed using GraphPad Prism 9.

### References:


1. Wang, W.X., Chen, W., Wu, C.Y., Zhang, C., Feng, J.J., Liu, P.X., Hu, Y.R., Li, H.Y., Sun, F., Jiang, K.*, et al.* (2023). Hydrogel-based molecular tension fluorescence microscopy for investigating receptor-mediated rigidity sensing. Nat. Methods *20*, 1780-1789.
2. Woodside, M.T., Behnke-Parks, W.M., Larizadeh, K., Travers, K., Herschlag, D., and Block, S.M. (2006). Nanomechanical measurements of the sequence-dependent folding landscapes of single nucleic acid hairpins. Proc. Natl. Acad. Sci. USA *103*, 6190-6195.
3. De Gennes, P.G. (2001). Maximum pull out force on DNA hybrids. Cr. Acad. Sci. Iv-

Phys. *2*, 1505-1508.

4. Hatch, K., Danilowicz, C., Coljee, V., and Prentiss, M. (2008). Demonstration that the shear force required to separate short double-stranded DNA does not increase significantly with sequence length for sequences longer than 25 base pairs. Phys. Rev. E Stat. Nonlin. Soft Matter Phys. *78*, 011920.
5. Mosayebi, M., Louis, A.A., Doye, J.P., and Ouldridge, T.E. (2015). Force-Induced Rupture of a DNA Duplex: From Fundamentals to Force Sensors. ACS Nano *9*, 11993-12003.
6. Li, H.Y., Zhang, C., Hu, Y.R., Liu, P.X., Sun, F., Chen, W., Zhang, X.H., Ma, J., Wang, W.X., Wang, L.*, et al.* (2021). A reversible shearing DNA probe for visualizing mechanically strong receptors in living cells. Nat. Cell Biol. *23*, 642-651.
7. Piella, J., Bastús, N.G., and Puntes, V. (2016). Size-controlled synthesis of sub-10-nanometer citrate-stabilized gold nanoparticles and related optical properties. Chem. Mater. *28*, 1066-1075.
8. Liu, Y., Blanchfield, L., Ma, V.P.Y., Andargachew, R., Galior, K., Liu, Z., Evavold, B., and Salaita, K. (2016). DNA-based nanoparticle tension sensors reveal that T-cell receptors transmit defined pN forces to their antigens for enhanced fidelity. Proc. Natl. Acad. Sci. USA *113*, 5610-5615.
9. Cavalcanti-Adam, E.A., Volberg, T., Micoulet, A., Kessler, H., Geiger, B., and Spatz, J.P. (2007). Cell spreading and focal adhesion dynamics are regulated by spacing of integrin ligands. Biophys. J. *92*, 2964-2974.
10. Oria, R., Wiegand, T., Escribano, J., Elosegui-Artola, A., Uriarte, J.J., Moreno-Pulido, C., Platzman, I., Delcanale, P., Albertazzi, L., Navajas, D.*, et al.* (2017). Force loading explains spatial sensing of ligands by cells. Nature *552*, 219-224.

**Table S1: Sequence of probes (5’-3’).**

| | |
|---|---|
| **5-10 pN** | |
| Strand A: A25H (5 pN) | 5’-CHCH-/iNH2C6dT/TAATATATAGTTTTTTTTCTATA-3’ |
| Strand B: P22A | 5’-PHO-TATTAATT/iNH2C6dT/TTACTACAATATA-3’ |
| Strand C: P33AQ | 5’-PHO-TAACAATGCTTTTGCATTGTTATATATTGTAG/iNH2C6dT/-BHQ2-3’ |
| **7-19 pN** | |
| Strand A: A25H (7 pN) | 5’-CHCH-/iNH2C6dT/AGATATATTAGTTTTTTTCTAATA-3’ |
| Strand B: P20A | 5’-PHO-TATCTATT/iNH2C6dT/TTACGCGCGCG-3’ |
| Strand C: P35AQ | 5’-PHO-CGCGCGCGCGCTTTTGCGCGCGCGCGCGCGCGCG/iNH2C6dT/ -BHQ2-3’ |
| **17-41 pN** | |
| Strand A: A21H | 5’-CHCH-/iNH2C6dT/GTCGCTCGGTGCTTTTGCAC-3’ |
| Strand B: P29A | 5’-PHO-CGAGCGACATT/iNH2C6dT/TTCGACGACAGACCACA-3’ |
| Strand C: P43AQ | 5‘-PHO-CAGCGAGCCAGCTTTTGCTGGCTCGCTG/iNH2C6dT/GTGGTCT GTCGTCG-BHQ2-3’ |
| **Bi-7 pN** | |
| Strand A: A25H (Bi-7 pN) | 5’-CHCH-/iNH2C6dT/AGATATATTAGTTTTTTTCTAATA-3’ |
| Strand B: P22A (Bi-7 pN) | 5’-PHO-TATTAATT/iNH2C6dT/TTACTACAATATA-3’ |
| Strand C: P33AQ (Bi-7 pN) | 5’-PHO-TAAGAATGCTTTTGCATTGTTATATATTGTAG/iNH2C6dT/-BHQ2-3’ |
| **Single hairpin (7 pN)** | |
| Strand A: A25H (7 pN) | 5’-CHCH-/iNH2C6dT/AGATATATTAGTTTTTTTCTAATA-3’ |
| Strand B: P11QS | 5’-PHO-TATCTATT/iBHQ2dT/TT-SSH-3’ |
| Complementary Strand (7 pN) | AAAAATAGATATATTAGAAAAAAACTAATATATCTA |
| **Single hairpin (17 pN)** | |
| Strand A: A21H | 5’-CHCH-/iNH2C6dT/GTCGCTCGGTGCTTTTGCAC-3’ |
| Strand B: P14QS | 5’-PHO-CGAGCGACATT/iBHQ2dT/TT-SSH-3’ |
| Complementary Strand (17 pN) | AAAAATGTCGCTCGGTGCAAAAGCACCGAGCGACA |

**Table S2: Calculated the unzipping force ($F_{1/2}$) of the hairpin during different conditions.**

| Length (nt) | Sequence (5'-3') | $\Delta G_{fold}$ (kJ/mol) | | $\Delta G_{stretch}$ (kJ/mol) | | Δx (nm) | $F_{1/2}$ (pN) | |
|---|---|---|---|---|---|---|---|---|
| | | a | b | a | b | | a | b |
| 31 | TAGATATATTAGTTTTTTTCTAA TATATCTA | 36.0 | 21.9 | 23.5 | 24.7 | 11.6 | 8.5 | 6.7 |
| 44 | ACGCGCGCGCGCGCGCGCG CTTTTGCGCGCGCGCGCGCG CGCGT | 179.5 | 150.6 | 33.4 | 35.1 | 17.4 | 20.4 | 17.8 |
| 30 | TGTCGCTCGGTGCTTTTGCAC CGAGCGACA | 85.7 | 68.2 | 22.8 | 23.9 | 11.2 | 16.1 | 13.7 |
| 31 | TTAATATATAGTTTTTTTTTCTAT ATATTAA | 26.5 | 13.4 | 23.5 | 24.7 | 11.6 | 7.1 | 5.4 |
| 44 | ACTACAATATATAACAATGCTTT TGCATTGTTATATATTGTAGT | 90.8 | 66.4 | 33.4 | 35.1 | 17.4 | 11.9 | 9.7 |

**a**: 22°C, PBS: 157 mM $Na^+$;

**b**: 37°C, DMEM medium: 155.3 mM $Na^+$, 0.8 mM $Mg^{2+}$.

**Table S3: Employed various models to calculate the shearing force required for rupture dsDNA.**

| Calibration value (pN) | Toy model (pN) | | de Genne, $N_{open}$=0 (pN) | de Genne, $N_{open}$=3.5 (pN) | de Genne, $N_{open}$=7 (pN) |
|---|---|---|---|---|---|
| | $\Delta G\tau_{obs}/\ \Delta G_{bp}$=7 | $\Delta G\tau_{obs}/\ \Delta G_{bp}$=4 | | | |
| 41.32±0.32 | 35.09 | 49.67 | 50.32 | 44.34 | 35.84 |

**Table S4: Summary of the thresholds of ForceChrono probes.**

| | **Probe 1** | **Probe 2** | **Probe 3** |
|---|---|---|---|
| **Calculated threshold (22°C in PBS)** | 7.1-11.9 pN | 8.5-20.4 pN | 16.1~44.3 pN |
| **Calculated threshold (37°C in DMEM)** | 5.4-9.7 pN | 6.7-17.8 pN | 13.7-44.3 pN |
| **Calibrated threshold using magnetic tweezer (22°C in PBS)** | **-** | 6.68-19.19 pN | 17.26-41.32 pN |

**Table S5: Calculated the unzipping force ($F_{1/2}$) of the second hairpin of Bi-7 pN probe.**

| Length (nt) | Sequence (5'-3') | $\Delta G_{fold}$ (kJ/mol) | | $\Delta G_{stretch}$ (kJ/mol) | | Δx (nm) | $F_{1/2}$ (pN) | |
|---|---|---|---|---|---|---|---|---|
| | | a | b | a | b | | a | b |
| 44 | ACTACATTATATAAGAATGCTTT TGCATTGTTATATATTGTAGT | 60.1 | 39.4 | 33.8 | 35.1 | 17.4 | 8.9 | 7.1 |

**a**: 22°C, PBS: 157 mM $Na^+$;

**b**: 37°C, DMEM medium: 155.3 mM $Na^+$, 0.8 mM $Mg^{2+}$.

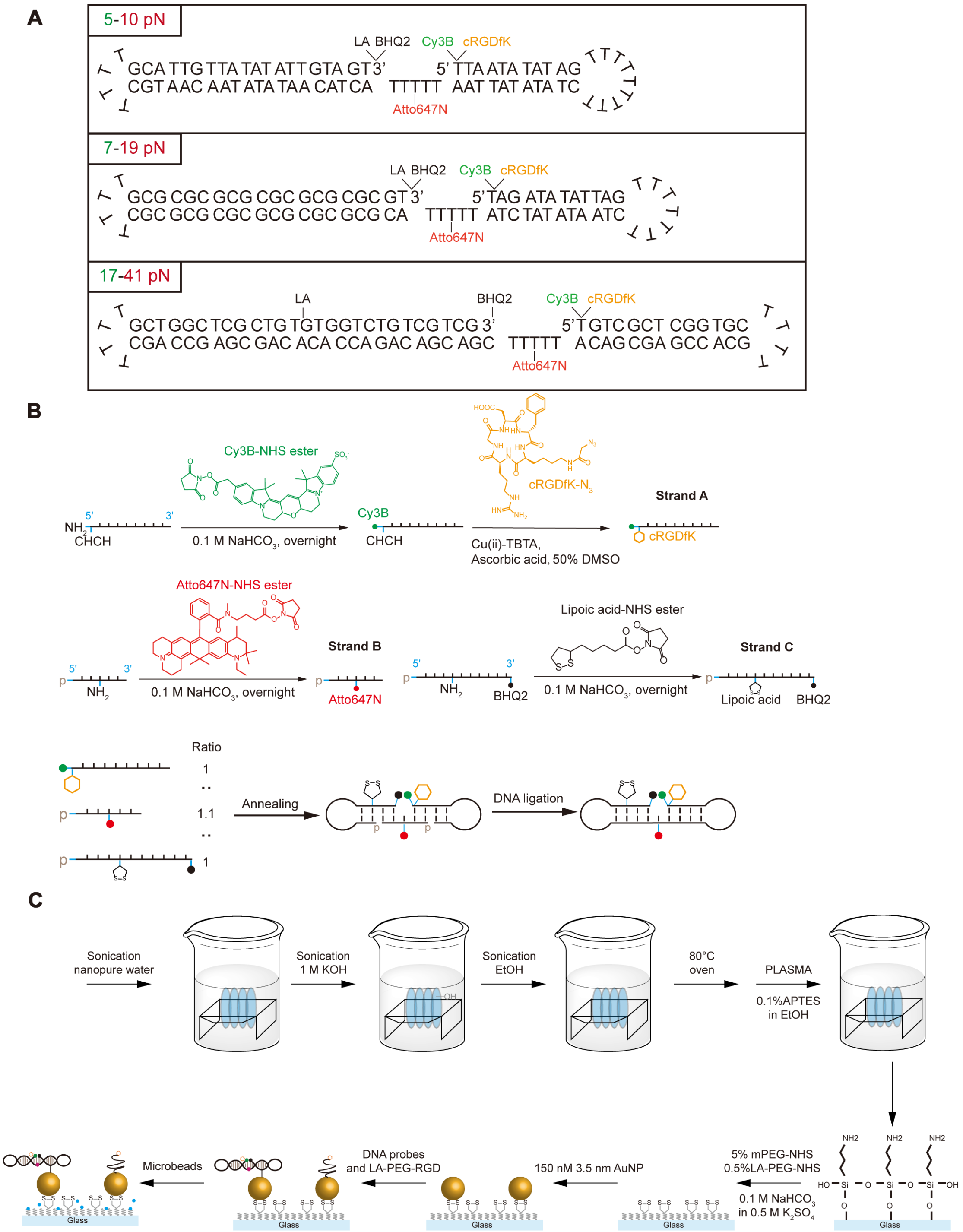


**Figure S1. Structure and synthesis scheme of ForceChrono probes. A,** Structures and oligonucleotide sequences of the 5-10 pN, 7-19 pN, and 17-41 pN ForceChrono probes. **B,** Step-by-step synthesis procedure of ForceChrono probes. **C,** Stepwise procedures for preparing ForceChrono probe-functionalized surfaces for single-molecule fluorescence imaging.

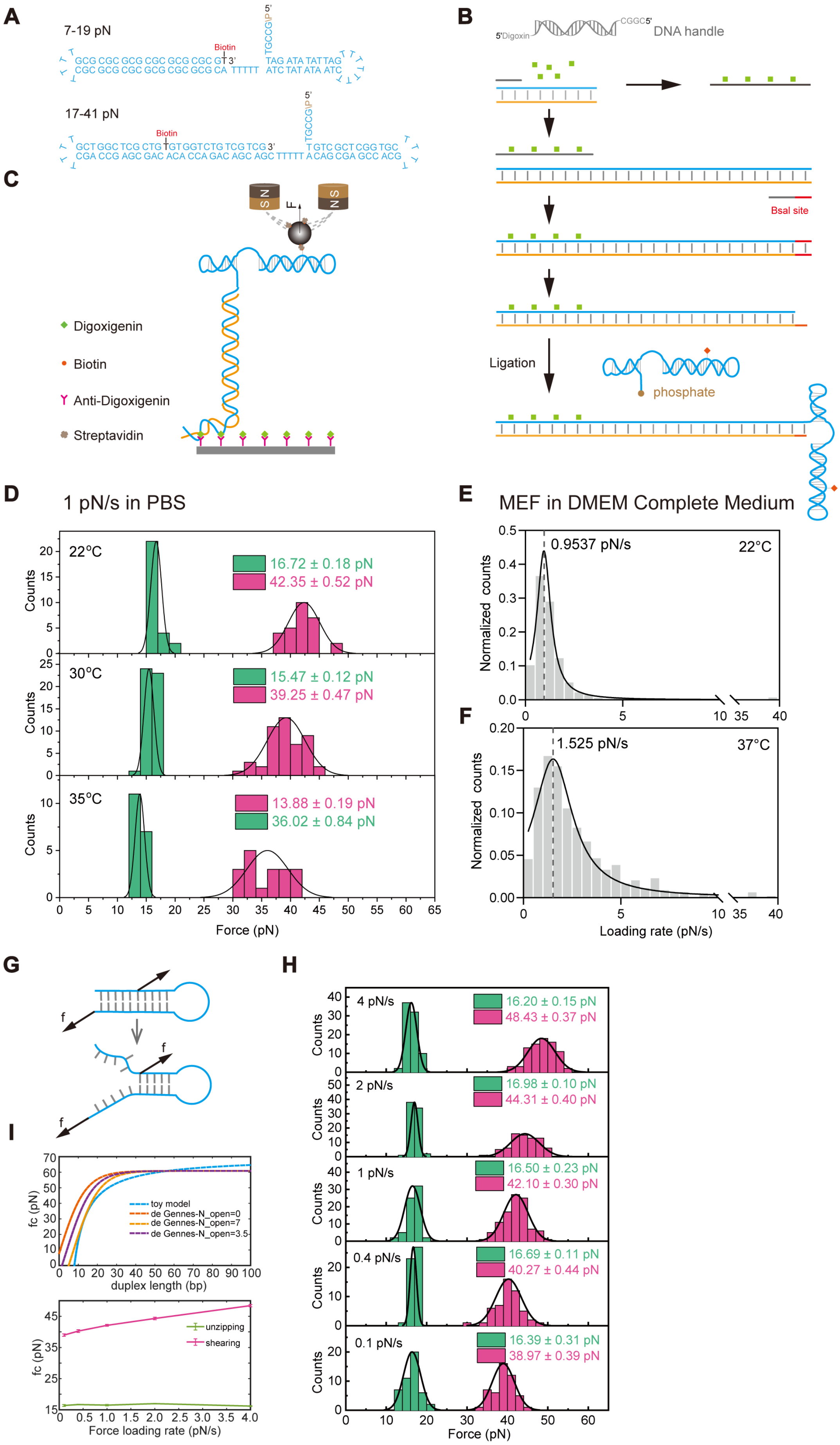

A
7-19 pN
Biotin
17-41 pN
B
5'Digoxin
DNA handle
BsaI site
Ligation
phosphate
C
Digoxigenin
Biotin
Anti-Digoxigenin
Streptavidin
D
1 pN/s in PBS
22°C
16.72 ± 0.18 pN
42.35 ± 0.52 pN
30°C
15.47 ± 0.12 pN
39.25 ± 0.47 pN
35°C
13.88 ± 0.19 pN
36.02 ± 0.84 pN
Counts
Force (pN)
E
MEF in DMEM Complete Medium
0.9537 pN/s
22°C
Normalized counts
F
1.525 pN/s
37°C
Normalized counts
Loading rate (pN/s)
G
H
4 pN/s
16.20 ± 0.15 pN
48.43 ± 0.37 pN
2 pN/s
16.98 ± 0.10 pN
44.31 ± 0.40 pN
1 pN/s
16.50 ± 0.23 pN
42.10 ± 0.30 pN
0.4 pN/s
16.69 ± 0.11 pN
40.27 ± 0.44 pN
0.1 pN/s
16.39 ± 0.31 pN
38.97 ± 0.39 pN
Counts
Force (pN)
I
fc (pN)
toy model
de Gennes-N_open=0
de Gennes-N_open=7
de Gennes-N_open=3.5
duplex length (bp)
unzipping
shearing
Force loading rate (pN/s)

**Figure S2. Calibration of ForceChrono probes using magnetic tweezers. A,** Geometries and composition of the 7-19 pN or 17-41 pN probes used for calibration. All probes have a 5' overhang with a GCCG sequence, and biotin is modified at the site where lipoic acid was previously modified to attach the probe to the magnetic bead. **B,** The production of connecting the 5' end of the probe to the multi-digoxigenin labeled DNA handle. **C,** Schematic depiction of the single-molecule magnetic tweezer experiments. The probes were immobilized to the anti-digoxigenin-coated surface through their DNA handles. Pulling forces were applied to a streptavidin-coated magnetic bead by permanent magnets to unfold the probe. **D,** Calibration of the 17-41pN probe at a loading rate of 1 pN/s in PBS at various temperatures. **E, F,** Determination of the loading rate of integrin on MEF cells at different temperatures using 17-41 pN probes. **G,** Several base pairs open at the end, and the number of these openings affects the probe's threshold (Upper). Various models have been employed to calculate the shearing force required for the rupture of double-stranded DNA (Lower). **H, I,** Calibration of the probe using a magnetic tweezer with different force loading rates.

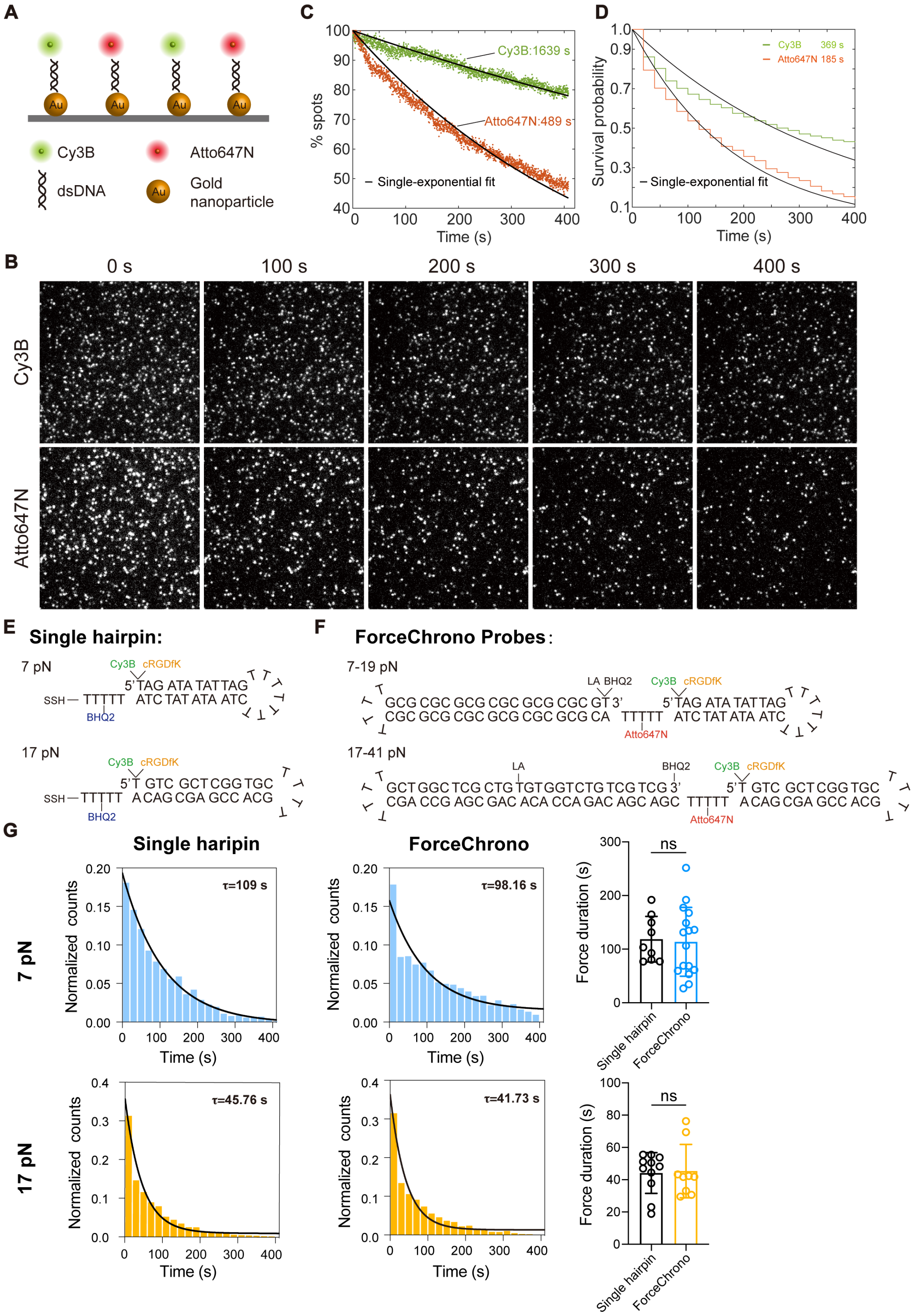


**Figure S3. Evaluating the stability and sensitivity of ForceChrono probes. A-D,** To investigate the impact of photobleaching and blinking of fluorescent dyes on the estimation of probe turn-on and turn-off events under force, we evaluated the stability of two fluorescent dyes, Cy3B and Atto647N, under experimental conditions (Non-phenol red DMEM complete medium supplemented with oxygen scavenging system and Trolox).

Cy3B and Atto647N were immobilized on an AuNP-modified surface via dsDNA in an equal ratio (**A**). We simultaneously excited Cy3B and Atto647N using 561 nm and 640 nm lasers with powers of 2.25 mW and 7.5 mW, respectively. The exposure time was set to 200 ms, and a continuous sequence of 2000 frames was captured for each channel (**B**). Evaluation of photobleaching by plotting the number of spots as a function of time and fitting the data using a single-exponential function. The photobleaching lifetime was determined from the exponential constant (**C**). Evaluation of blinking during excitation by tracking the intensity of single molecules and measuring the duration from the first frame to the first transition to a dark state or photobleaching. Survival analysis was used to determine the decay lifetime (**D**). **E,** Structures and oligonucleotide sequences of the 7 pN and 17 pN Single hairpin probes, with the sequence of the single hairpin identical to the first hairpin in the ForceChrono probes. **F,** Structures and oligonucleotide sequences of the 7-19 pN and 17-41 pN ForceChrono probes. **G,** The left graph displays the force duration distribution of integrins across all MEF cells, with the mean duration represented by black lines fitted using an exponential function. The right graph illustrates the average force duration measured by each MEF cell using different probes. Each point represents the mean duration of a cell. The error bars represent mean ± s.d. from three independent experiments. Two-tailed Student's *t*-tests are used to assess statistical significance.

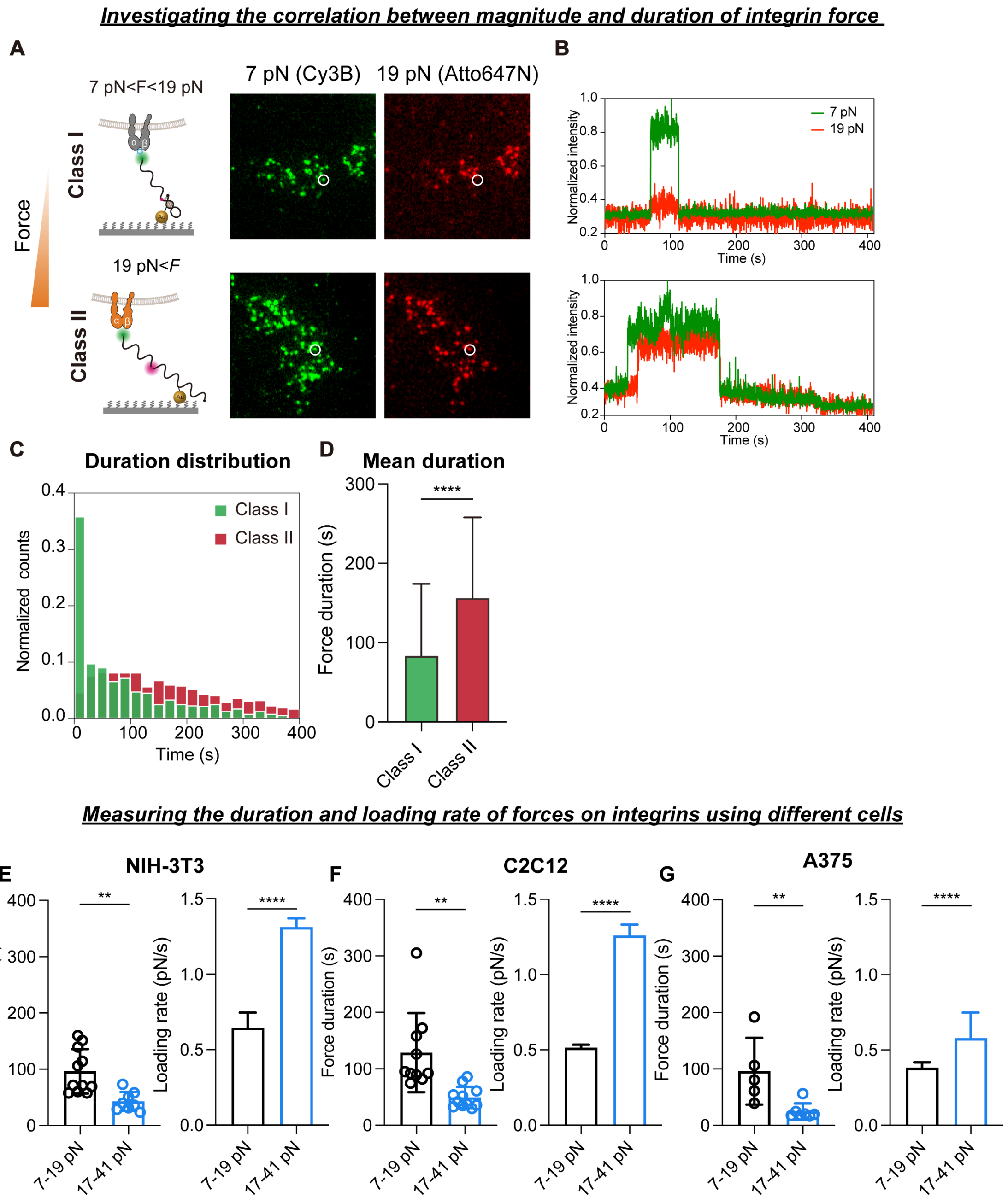


**Figure S4. Investigation of the mechanical force duration and loading rate on integrins using ForceChrono probes. A,** Left: schematic representation illustrating the division of integrins into two classes based on force magnitude. Right: representative molecular force signal reported by 7-19 pN probes. **B,** Representative fluorescence intensity traces for the two classes of integrins. **C,** Distribution analysis of the duration of mechanical forces exerted on the two classes of integrins. **D,** Since the force duration of class II integrins cannot be fitted with an exponential function, the mean duration of a single integrin is shown here, based on data from over 1000 integrins. **E-G,** Measurement of the average duration and loading rate of force on integrins using 7-19 pN or 17-41 pN ForceChrono probes in NIH-3T3 (**E**), C2C12 (**F**) and A375 (**G**) cells. For force duration, each point represents the mean force duration of a cell. The error bars represent mean ± s.d. from three independent experiments. For loading rate, the error bars represent mean

± 95% CI from three independent experiments. Two-tailed Student's *t*-tests are used to assess statistical significance.

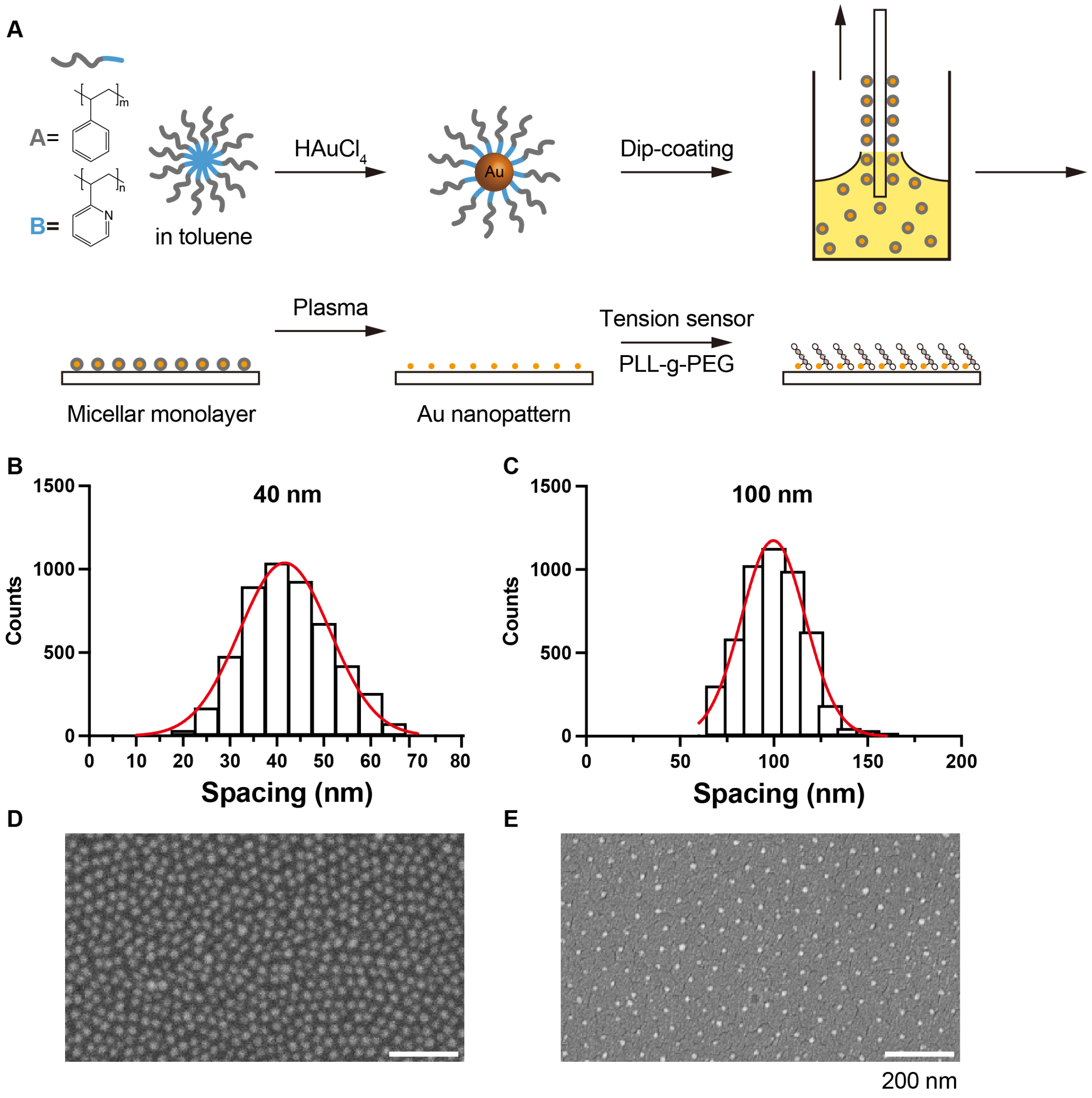


**Figure S5. Characterization of nanopatterned AuNP arrays. A,** Fabrication of a gold nanoparticle nanopatterned substrate utilizing block copolymer micelle nanolithography and functionalization with ForceChrono probes. **B, C,** The spacing of BCMN-generated nanopatterned AuNP arrays. **D, E,** Scanning electron micrograph of gold nanoparticle nanopatterned substrate. Scale bar=200 nm.

**Movie captions:**

**Movie S1:** Single-molecule fluorescence imaging results of mechanical force signal of MEF cells on 17-41 pN probe. Cy3B and Atto647N signals were collected simultaneously to image 2000 frames without intervals at 200 ms exposure time.

**Movie S2:** Single-molecule fluorescence imaging results of mechanical force signal of MEF cells on 5-10 pN probe. Cy3B and Atto647N signals were collected simultaneously to image 2000 frames without intervals at 200 ms exposure time.

**Movie S3:** Single-molecule fluorescence imaging results of mechanical force signal of *Vcl* KO MEF cells on 5-10 pN probe. Cy3B and Atto647N signals were collected simultaneously to image 2000 frames without intervals at 200 ms exposure time.
